# Detect Genomic G-Quadruplexes in Living Animal Cells with a Tiny Artificial Protein Probe

**DOI:** 10.1101/2020.05.06.080622

**Authors:** Ke-wei Zheng, Jia-yu Zhang, Yi-de He, Jia-yuan Gong, Cui-jiao Wen, Juan-nan Chen, Yu-hua Hao, Zheng Tan

## Abstract

G-quadruplex (G4) structures formed by guanine-rich nucleic acids are implicated in essential physiological processes and serve as important drug targets. The genome-wide detection of G4s in living cells is important for exploring the biological role of G4s but has not yet been achieved due to the lack of a suitable G4 probe. We engineered a 6.7 kDa G4 probe (G4P) protein that binds G4s with high affinity and specificity. We used it to capture G4s in living human, mouse, and chicken cells with the ChIP-Seq technology, yielding genome-wide landscape as well as details on the positions, frequencies, and sequence identities of G4 formation in these cells. Our results indicate that transcription is accompanied by a robust formation of G4s in genes. In human cells, we detected up to >123,000 G4 peaks, of which >1/3 had a fold increase of ≥5 and were present in >60% promoters and ~70% genes. Being much smaller than a scFv antibody (27 kDa) or even a nanobody (12-15 kDa), we expect that the G4P may find diverse applications in biology, medicine, and molecular devices as a G4 affinity agent.

## INTRODUCTION

G-quadruplexes (G4s) are four-stranded secondary structures formed by guanine-rich nucleic acids. Putative G-quadruplex forming sequences (PQSs) are abundant in the genomes of animal cells with particular enrichment near transcription start sites (TSSs), implying an essential role of G4s in transcription (1). Accordingly, they are emerging as a new class of drug targets for pharmaceutical applications. As such, the detection and quantitation of G4s in genomes with sequence identity are indispensable for exploring the biological function and drug targeting of G4s. Recently, G4s have been detected in chemically fixed human cells by immunostaining (2) or immunoprecipitation (3,4) with a G4 antibody. In these applications, the binding of antibodies to the G4s occurs in a non-native cellular environment after a series of treatments. It is not known how G4s would be reserved or affected during the permeabilization, staining, and fragmentation, or DNA purification (5–7) in the duplex genome DNAs in which the hybridization of two complementary DNA strands competes against the formation of G4s. A recent study has shown that G4s formed in a duplex DNA in transcription collapse fast when the RNA transcript annealed with the DNA template is removed (8) even in an environment in which DNA hybridization is significantly weakened and G4 simultaneously stabilized (9). For these reasons, G4 identification in living cells is desired as an alternative option for the G4 community.

Unfortunately, antibodies are not suitable for living cells because they do not permeate into cells. Besides, the reductive environment of the cytoplasm of living cells is not compatible with the formation of the disulfide bonds required for maintaining the tertiary structure of antibodies. An attempt to use an antibody in living human cells detected G4s mostly in the telomeres in which long clusters of PQS are present (10). On the other hand, native G4-interacting proteins are not appropriate either in that they usually possess multi-functional domains such that they may interact with other proteins or macromolecules besides the G4s. There are chances for them to be brought to targets indirectly or subject to complex interactions, resulting in non-specificity or impeded recognition.

To overcome these difficulties, we engineered a 64 amino acids (64-aa) 6.7 kDa protein based on the G4-binding domain of the RHAU, an RNA helicase able to bind and unwind G4s (11). This G4 probe (G4P) comprises only two small G4-binding domains linked by a flexible and optimized linker in between. Such a simple composition minimizes non-specific interactions with other proteins. Moreover, the synergy between two binding domains dramatically improves affinity and selectivity towards G4s. Expression of the G4P in cells followed by chromatin immunoprecipitation (termed G4P-ChIP) allowed us to capture G4s in living human, mouse, and chicken cells through the ChIP-Seq technology, revealing genome-wide landscape and details on the locations, frequencies, and sequence identities of G4 formation in these cells.

## MATERIALS AND METHODS

### ChIP-Seq data analysis

Clean paired-end sequencing data in fastq format were mapped to the human genome (hg19 or hg38 when comparing with downloaded hg38 data) using the Bowtie2 software (12) with the sensitive-local preset and --no-unal, --no-discordant, --no-mixed parameters. Mapped reads were written to bam files after being filtered by the samtools view (13) to remove low-quality alignments with the parameter −q 20 and by samtools rmdup to remove duplicates. Reads enrichment was calculated using the deeptools (14) plotEnrichment with the bam files. Reads bam files were also processed by the deeptools bamCompare to produce bigwig coverage files in subtract or ratio mode and normalized to RPKM. Profiles and heatmaps of reads were generated from the bigwig files using the deeptools computeMatrix followed by plotProfile and plotHeatmap, respectively, with region bed files derived from the NCBI RefSeq bed file downloaded from the UCSC website (http://genome.ucsc.edu/) unless otherwise indicated. Coordinate duplicates in the bed files were removed. Peaks of reads enrichment were identified with the macs2 software (15) using --qvalue 0.001, --keep-dup 1, and default values for the other parameters. ChIP-Seq data from public repositories were downloaded from the GEO (https://www.ncbi.nlm.nih.gov/geo/) or Encode (https://www.encodeproject.org/) database and processed as described above whenever applicable. Original sequencing (fastq) and processed (narrowPeak, bigwig) G4P-ChIP datasets have been deposited in and can be downloaded from the NCBI Gene Expression Omnibus (GEO) under accession code GSE133379.

Detailed materials and methods are in Supporting Information.

## RESULTS

### G4P binds canonical and non-canonical G4s *in vitro*

Our G4P core is a 64-aa artificially engineered protein (**Figure 1A**) composed of two identical 23-aa G4-binding domains (RHAU23) from the RHAU protein that has a preference to bind parallel G4s with a dissociation constant (*K*_d_) of ~1 μM (11). The RHAU23 contained a 13-aa RHAU-specific motif that functions as a major determinant for the affinity and specificity toward G4s in the original RHAU (16). According to the solution structure of the RHAU23-G4 complex, each of the two terminal guanine quartets (G-quartets) of a G4 can bind an RHAU23 (11). Therefore, the two RHAU23 units in the G4P are expected to clamp onto the two terminal-G-quartets of a G4, reinforcing the binding through a synergy between two binding domains (**Figure 1B**). We added a HIS and a 3xFLAG tag at the N- and C-terminal of the G4P core, respectively (**Figure 1A**). This 113-aa G4P was expressed in *E. coli* and purified by affinity chromatography using the HIS tag.

**Figure 1.**
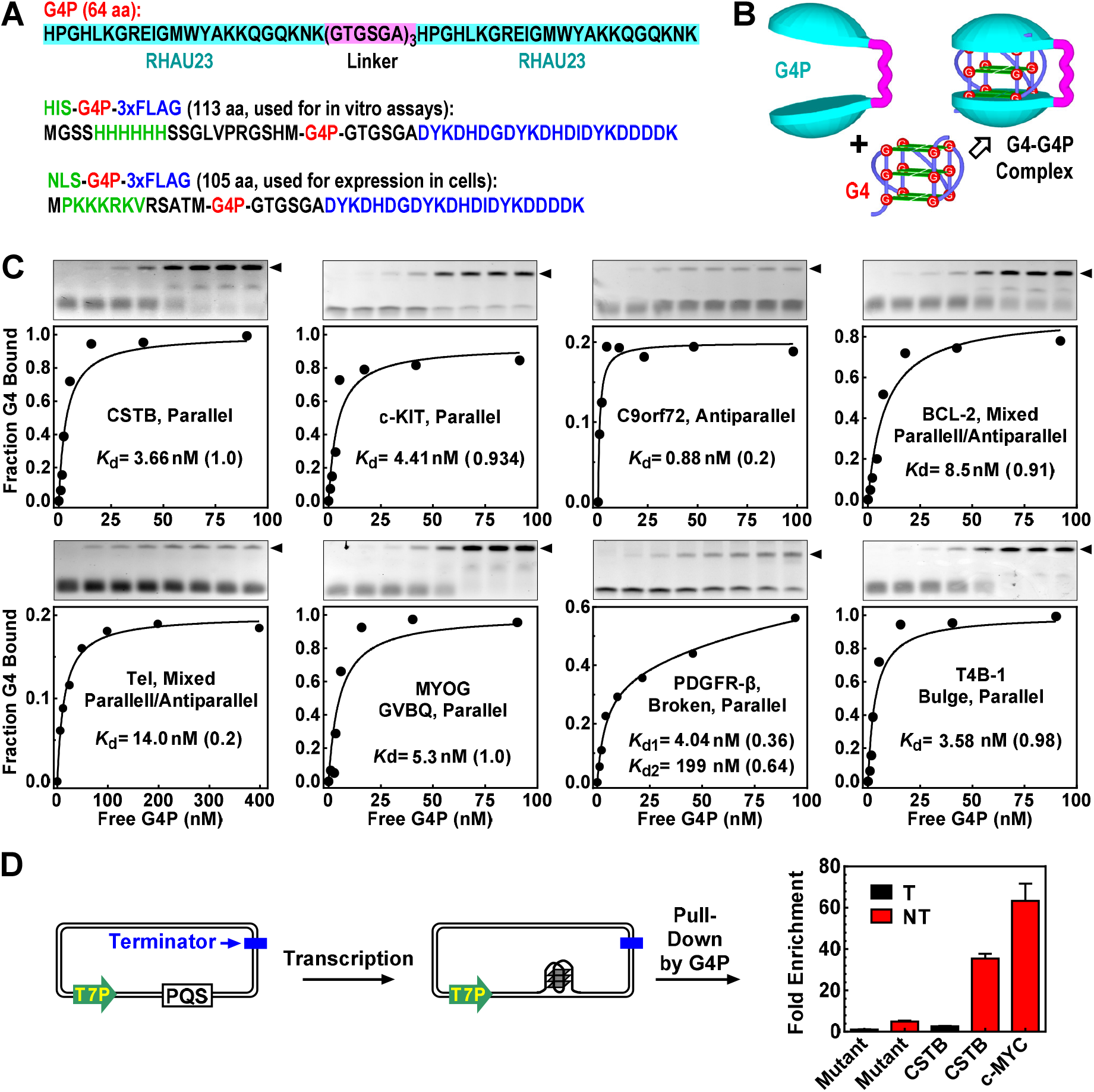
G4P binds G4s in vitro. (A) The amino acid sequence of G4P. The one with a nuclear localization sequence (NLS) was used for detecting G4s in cells. The one with a HIS tag was for in vitro assays. A 3xFLAG at the C-terminal was for binding with the anti-FLAG antibody. (B) Anticipated clamping-binding of a G4 by a G4P. (C) Dissociation constant *K*_d_ of G4P to G4s of different folding conformation determined by EMSA. *K*_d_ values were obtained by fitting the data of each DNA (**Table S1**) to a model assuming the existence of two subpopulations of G4s. The number in parenthesis indicates the fraction of the G4 subpopulation associated with the corresponding *K*_d_. One *K*_d_ is given when the fitting produced two identical *K*_d_ values or the other is >1 M. (D) Pull-down of transcriptionally generated G4 in plasmids by G4P. A PQS or mutant motif (**Table S2**) was placed on either the non-template (NT) or template (T) strand downstream of a T7 promoter (T7P) in a plasmid. The plasmid transcribed with T7 RNA polymerase was incubated with G4P. G4P-G4 complex was then pulled-down in the presence of an internal reference plasmid using anti-FLAG beads and G4-bearing plasmid was quantitated by qPCR relative to the internal plasmid. Fold enrichment was normalized to the mutant motif on the template strand.

Besides the canonical G4s, there are three types of well-characterized non-canonical G4s, i.e. G4s with one loop of 8-15 nucleotides (4GL15) (17), G-vacancy-bearing G4s (GVBQ) (18), G4s with a bulge of one non-G nucleotide (Bulge) (19), respectively. We first assessed the binding activity of the G4P to several representative G4s (**Table S1**) by the electrophoretic mobility shift assay (EMSA) (11). For all the DNAs tested, effective mobility shift was observed above the original DNA band in positive correlation to G4P concentration (**Figure 1C** and **Figure S1A**, arrowhead, gels), which is in contrast to the RHAU23 that largely failed to bind the G4s (**Figure S1B**). The tags did not affect the binding of G4P to G4s (**Figure S1C**). Similar to the RHAU23, the G4P did not bind the non-G4 DNAs, including i-motif sequences, single-stranded DNA (ssDNA), and hairpin double-stranded DNA (dsDNA) (**Figure S1D**).

In some cases, we noticed that the G4P did not bind all the DNAs in a sample even at saturating concentration (**Figure 1C** and **Figure S1A**). For example, G4P bound only 20% of the C9orf72 and Tel DNAs. Because these DNAs have been reported to adopt an antiparallel (20) and mixed parallel/antiparallel (21) conformation, respectively, that is not favored by the original RHAU23 (11), there was a possibility that the 20% of the DNAs instead adopted a conformation recognized by the G4P. These results indicated that the conformations for a given DNA might be heterogeneous and a fraction of the G4s could still be recognized by the G4P. Therefore, we fitted the EMSA data to a two-population binding model for simplicity to derive the dissociation constant *K*_d_. High-affinity *K*_d_ at low-nM was obtained for each PQS with various fractional quantities (**Figure 1C, Figure S1A**). The results also suggested that G4P could bind more than one folding conformations with distinctive affinities as demonstrated by the PDGFR-β G4s that exhibited two distinct *K*_d_ values.

We next examined G4 binding under a more physiologically relevant condition to pull down a plasmid containing a G4 induced by transcription (**Figure 1D, Table S2**). The plasmid accommodated a CSTB, c-MYC, or a mutated motif on the non-template or the template strand. Transcription with T7 RNA polymerase efficiently induces a formation of G4 on the non-template but marginally on the template strand (22). Accordingly, the G4s on the non-template strand led to a high enrichment of the corresponding plasmid. A marginal enrichment of the plasmid with the mutant motif on the non-template strand might be attributed to the possible formation of a weak DNA:RNA hybrid G4 involving the G_2_ and G_4_ tracts from the DNA and RNA transcript (23,24).

The formation of the G4 in the plasmid was unlikely induced by the G4P because G4 was not detected when the CSTB motif was placed on the template strand, which we previously described as a strand-biased G4 formation in transcription (22). Furthermore, G4 was not detected by circular dichroism (CD) spectroscopy (**Figure 2A**) and EMSA (**Figure 2B**, lanes 3 and 5) when a CSTB-containing dsDNA (**Table S3**) was incubated with G4P. Instead, the G4P only captured the G4s that already existed (**Figure 2B**, lanes 7 and 9 versus 3 and 5). When we incubated G4P with single-stranded PQS motifs in a Li^+^ solution, a condition that does not support G4 formation, the CD spectra of the DNAs remained virtually unchanged (**Figure 2C**, dashed curves). Notably, the G4s formed in a K^+^ solution maintained their folding topology after they were bound by G4Ps (**Figure 2C**, solid curves), which ensured that the recognition remained even in the G4P-G4 complexes.

**Figure 2.**
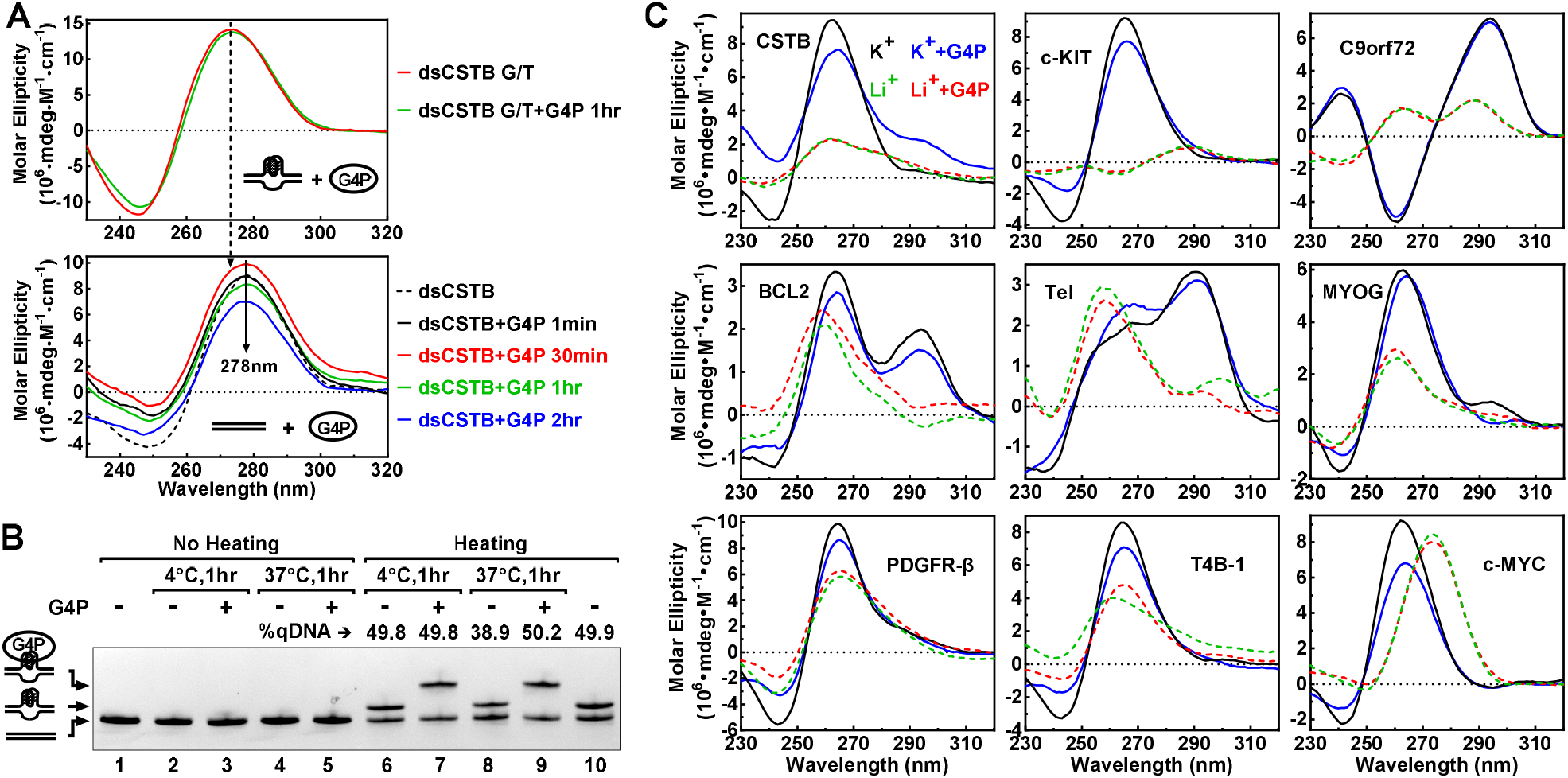
G4P did not induce a formation of G4 in dsDNA. (A) Circular dichroism (CD) spectra of a CSTB PQS-containing dsDNA in the absence and presence of G4P at 1:1 molar ratio to DNA. The DNA strands in the top panel were not complementary to each other within the PQS region to enable G4 formation. The DNA strands (**Table S3**) in the bottom panel were fully complementary to each other. They were incubated with or without G4P for the indicated time at RT. (B) The dsDNA used was heated to generate G4 and then incubated without or with G4P at 4 or 37 °C, respectively, for 1 hour before EMSA. At 4 °C, the percentage of G4-bearing DNA (qDNA) remained unchanged when the DNA was left free or bound by G4P (lanes 6 and 7 versus 10). At 37 °C, the percentage of qDNA decreased by ~20% when the DNA was left free but remain unchanged when bound by G4P (lanes 8 and 9 versus 10). (C) CD spectra of G4 structures in the absence or presence of G4P. DNAs (**Table S1**) were incubated without or with G4P at 1:1 molar ratio at RT for one hour in 150 mM Li^+^ or K^+^ solution. G4s are stabilized by K^+^ but not by Li^+^.

### G4P recognizes G4s in living cells

To detect G4s in living cells, we added a nuclear localization sequence (NLS) and a 3xFLAG tag at the N- and C-terminal of the G4P core, respectively, without compromising the binding to G4s (**Figure S1C**). We then expressed this 105-aa G4P in cultured human A549 cells by transfection with a plasmid and performed ChIP-Seq on the G4P-G4 complex (**Figure 3A**). The G4P reads were mapped to the human genome using the Bowtie2 (12) and high-quality reads (−q=20) were subsequently obtained using the Samtools (13) and then their distribution on genome analyzed with the Deeptools (14). The binding of the G4P to G4s was first demonstrated by the enrichment of G4P reads mapped to the canonical PQS motifs defined by a consensus of G_≥3_(N_1-7_G_≥3_)_≥3_ (24,25) (**Figure 3B**). We term this type of PQSs 4G PQSs to distinguish them from the non-canonical ones. The enrichment disappeared when the coordinates of the 4G PQSs were shuffled to random 4G PQS-free locations using the Bedtools ShuffleBed (26) and the reads around these non-PQS regions counted. Since PQSs are concentrated around TSSs where DNA is more accessible to fragmentation than other regions, the input reads at real 4G PQSs was greater than at the shuffled. The binding of G4P to G4s was further illustrated by a peak when the G4P reads were profiled around the center of the 4G PQSs, which appeared only in the G4P-ChIP but not so in the input sample (**Figure 3C**). The tiny peak in the input could be attributed to a greater openness of the binding site than the other regions, as “features of open chromatin” in ChIP-Seq (27). Similarly, shuffling the coordinates of the 4G PQSs also removed the difference between the two samples. As a routine independent validation, we observed an enrichment of G4P at the 4G PQSs by ChIP-qPCR (**Figure S2**).

**Figure 3.**
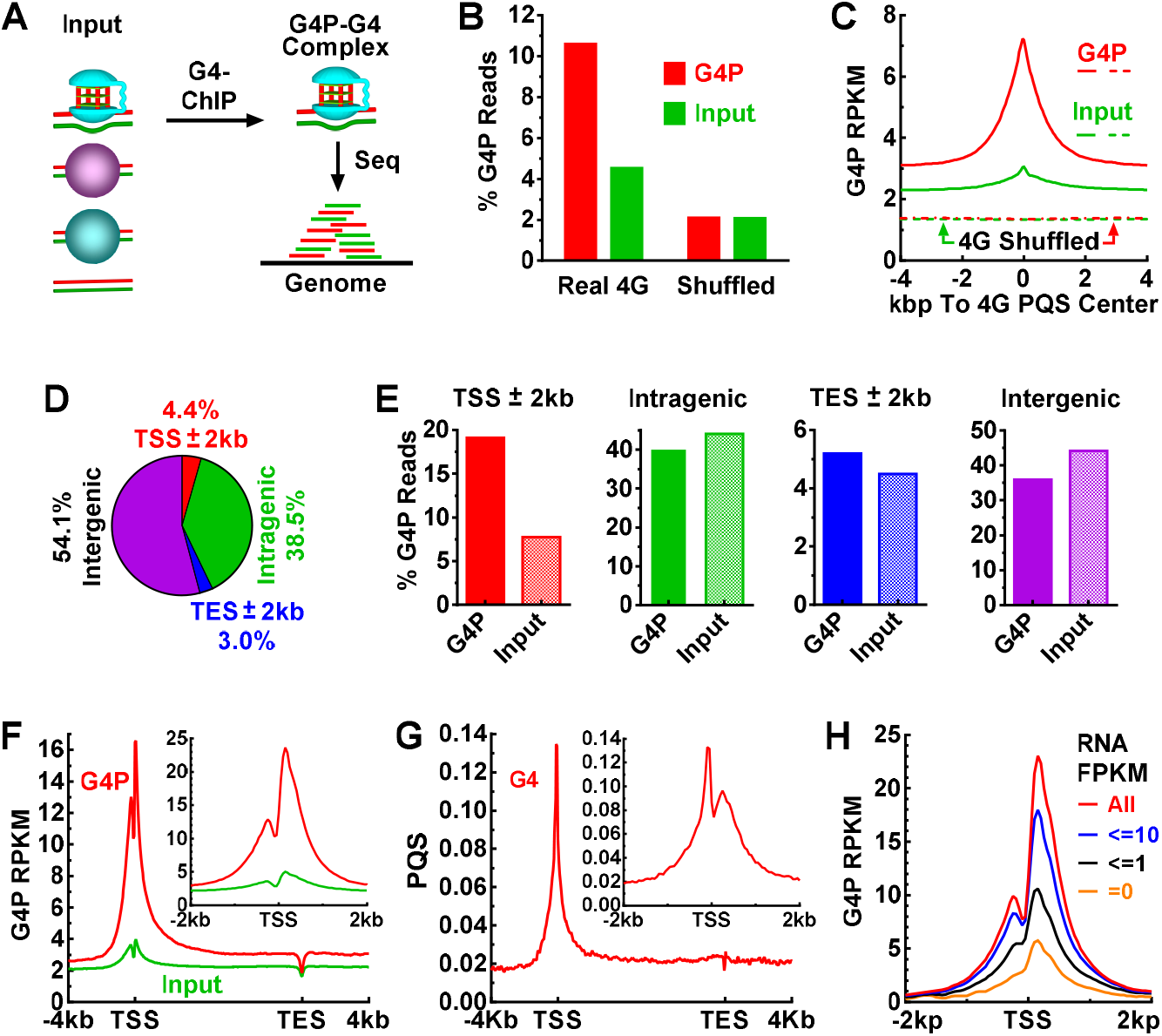
G4P binds genomic G4s in human A549 cells. (A) G4P-G4 complex was enriched from fragmented input DNA by chromatin immunoprecipitation (G4-ChIP) and then subjected to sequencing (Seq) to identify G4 formation in the genome. (B) G4P reads at 4G PQSs as a percent of the total reads mapped to the genome. (C) Enrichment of G4P at 4G PQSs. (D) Percent nucleotides and (E) percent G4P reads in genomic regions. (F) G4P reads and (G) PQSs distribution across RefSeq genes and around TSSs (insert). (H) G4P reads distribution around TSS in genes of different RNA expression levels. The signal in (F) is normalized to RPKM and G4P signal in (H) is expressed as (G4P Reads) - (Input Reads) and normalized to RPKM.

To find out where G4 forms in the genome, we surveyed G4 formation in the four functional regions, i.e. TSS±2kb (promoter), intragenic, TES±2kb, and intergenic regions (**Figure 3D**), where TSS and TES represented transcription start and end site, respectively. The G4P in the TSS±2kb region counted >19% of the total mapped reads, more than twice the input in this region, which was in contrast to the other three regions (**Figure 3E**). This result indicated that the gene promoters were hot sites of G4 formation in the human genome. We further profiled G4P reads distribution in the RefSeq genes and found that G4s were concentrated at both sides of TSSs, with two peaks merging at the TSSs (**Figure 3F**). The distribution largely correlated with the presence of 4G PQSs across the TSSs (**Figure 3G**) and agreed with our previous *in vitro* studies in which transcription efficiently induced G4s at both sides of a TSS (18,22–24,28–30).

The distribution of the two features between TSS and TES were distorted due to the scaling on the gene bodies of different sizes. A better correlation was seen when the profiles were plotted across TSS only (**Figure 3**, F and G, inserts). It is noted that the extent of G4 formation was smaller at the upstream side than at the downstream side of the TSSs while the occurrence of PQS was the opposite. This phenomenon could be explained by the asymmetrical transcription activities crossing TSS. For example, synthesis of RNA occurs at the downstream side, generating an RNA:DNA hybrid structure known as R-loop (31), which has been shown to reinforce transcriptional G4 formation by suppressing the hybridization of the duplex DNA (8). *In vitro* transcriptional formation of G4s positively correlates with transcription activity (8,32). To find out the *in vivo* situation, we grouped TSSs according to their expression level and found greater G4 formation was associated with higher gene expression (**Figure 3H**).

### G4P detects non-canonical G4s

As a natural deduction, more PQS motifs should lead to more G4 formation in a region. We thus plotted G4P profiles as a function of the number of 4G PQSs in the TSS±3kb regions. The reads density decreased as we gradually excluded the regions with a greater number of 4G PQS (**Figure 4A**). However, when all the 4G PQS-positive regions were excluded, the peak only dropped by ~60% in the remaining ~1/3 of the TSSs. This fact suggested that the canonical 4G PQSs did not account for all the G4s recognized by the G4P and other forms of G4s were also captured. We, therefore, searched for PQSs of 4GL15 (17), GVBQ (18), Bulge (19), respectively (**Figure 4B**). These PQSs were similarly concentrated near TSSs as the 4G PQSs did (**Figure 4C**), suggesting that they were also evolutionally selected to form G4s and function in cells. Hence, they might all be targets of the G4P in cells as they were in *vitro* (**Figure 1C** and **Figure S1A**).

**Figure 4.**
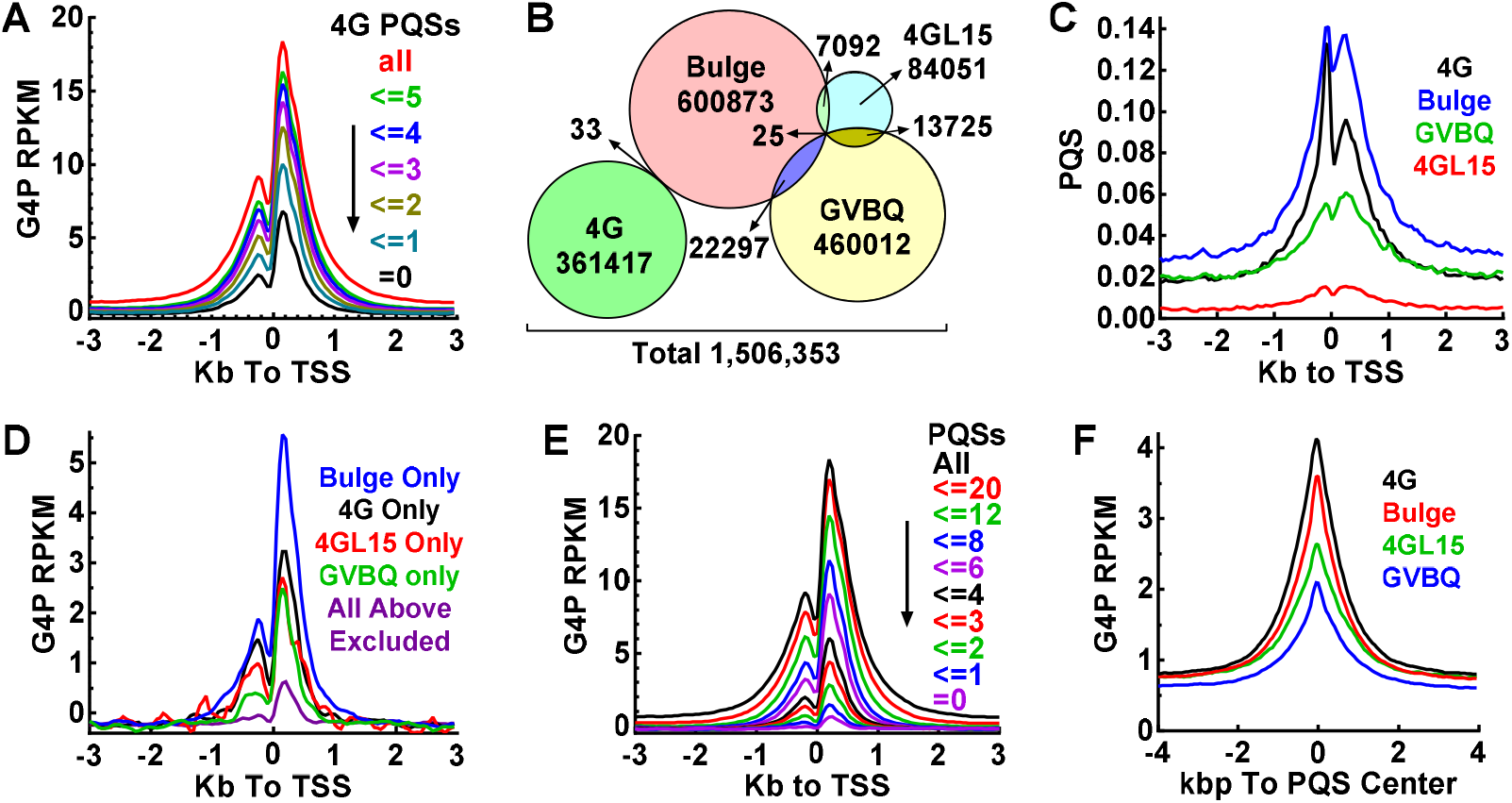
G4P binds canonical and non-canonical genomic G4s in human A549 cells. (A) Enrichment of G4P depends on the 4G PQS load. (B) Amount of four subtypes of PQSs in the human genome. Overlaps show motifs carrying more than one subtype of PQSs. (C) Distribution of 4G, 4GL15, GVBQ, and Bulge PQSs around TSSs. (D) Distribution of G4P reads around TSSs with only the indicated subtype of PQS. (E) Enrichment of G4P depends on the load of four types of PQS. (F) Enrichment of G4P at subtypes of PQSs. G4P signal is expressed as (G4P Reads) - (Input Reads) and normalized to RPKM. Numbers of PQS motifs within the TSS±3kb region are given in panels A and E, respectively.

To verify the detection of non-canonical G4s, we filtered out those TSSs carrying more than one subtype of PQSs in the TSS±3kb region and examined the contribution of each subtype. The result (**Figure 4D**) shows that G4P recognized each of them. When the four subtypes of PQSs were all excluded, the reads peak decreased to a very low level (**Figure 4D**, purple curve) that might represent recognition of other rare types of PQSs, for example, those with a loop larger than 15 nucleotides (17) or a bulge of more than one nucleotide (19). When the number of all the four subtypes of PQSs were counted, a positive correlation was also obtained between G4 formation and PQS load (**Figure 4E**). The enrichment of G4P at each subtype of PQS suggested that the ability of G4 formation followed the order of 4G>Bulge>4G15L>GVBQ (**Figure 4F**).

### Overview and statistics of G4 formation in living cells

To overview the landscape of G4 formation in human genes, we generated a 3D heatmap of G4P around TSSs for the A549 cells. It showed that G4s formed in most of the genes near TSSs (**Figure 5A**) in the presence of PQSs (**Figure 5B**). A tiny fraction of TSSs showed an obvious enrichment of PQSs (**Figure 5B**, blue arrowhead) but with little G4P (**Figure 5A**), indicating an absence or low probability of G4 formation in a small fraction of the PQS motifs. We also expressed the G4P and detected G4s in cultured human NCI-H1975, HeLa-S3, 293T, mouse 3T3, and chicken DF-1 cells, respectively. G4 formation in these cell lines all displayed similar landscapes crossing TSSs (**Figure S3**) and genes (**Figure S4**) although their magnitudes varied (**Figure S4**).

**Figure 5.**
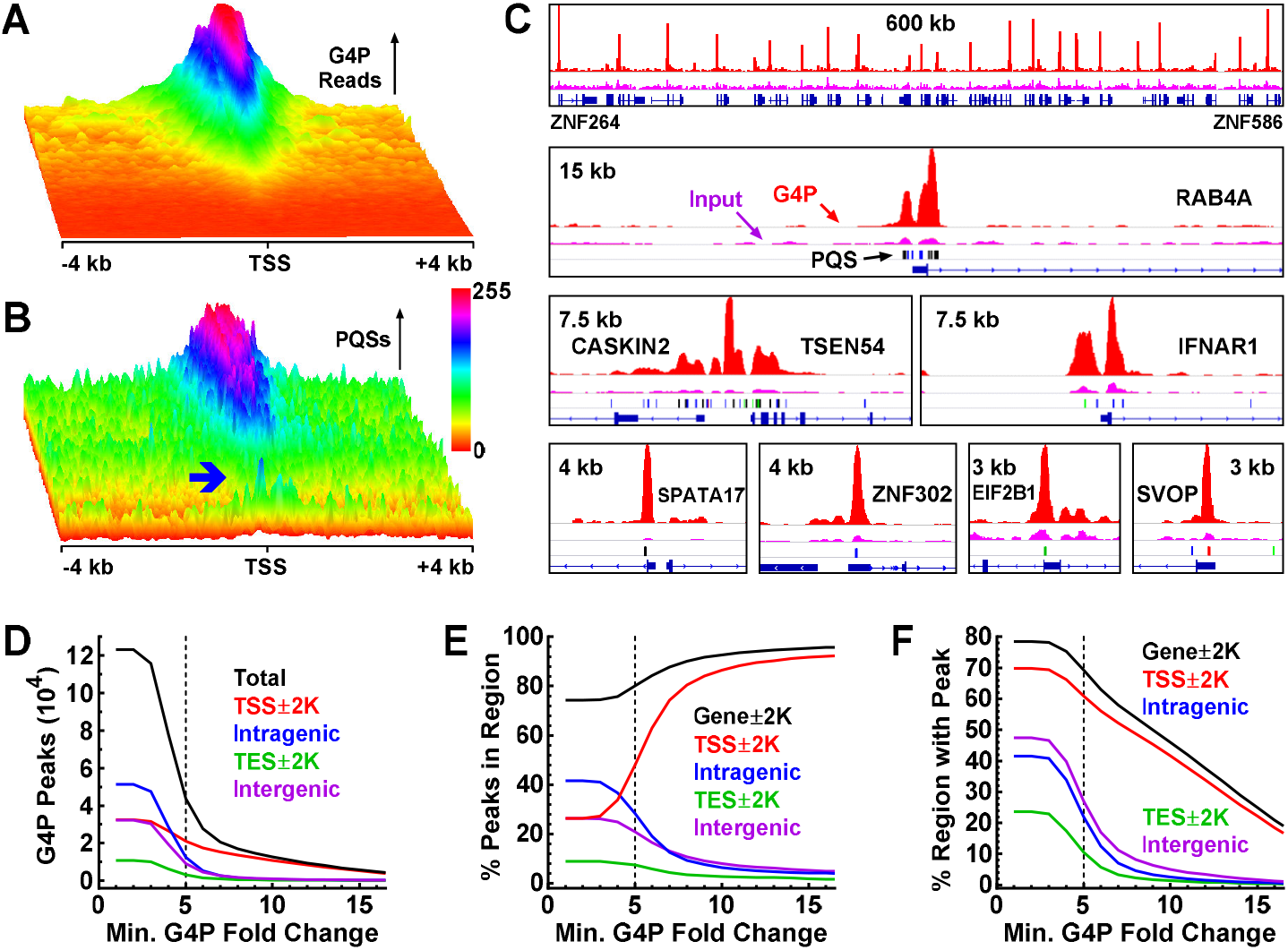
Overview and statistics of DNA G4 formation in human A549 cells. (A) G4P enrichment and (B) the presence of PQSs in the TSS±4kb regions of RefSeq genes showing G4 formed at where PQS existed. The two heatmaps were produced over the same TSS region file sorted in descending order on the maximum of G4P reads. The blue arrowhead shows a small fraction of PQS motifs with a low probability of G4 formation. Values in each heatmap were normalized independently to the range of 0-255 to enhance the visual resolution. (C) Examples of G4 formation depicted by G4P peaks. Reads signals for G4P and input are in red and magenta, respectively. Color bars beneath the input lane indicate 4G (black), 4GL15 (red), Bulge (blue), and GVBQ (green) motifs, respectively. Genomic range and genes are indicated in each panel. (D-F) Statistics of G4 formation as a function of G4P peak fold change.

To identify individual G4-binding events, we performed peak calling on the G4P reads from the A549 cells using the MACS2 (15), which identified 123,274 G4P peaks indicative of G4-G4P interactions. In **Figure 5C**, we present examples of G4 formation indicated by G4P peaks in association with the presence of different types of PQSs. More examples from other cell lines are given in **Figure S5–S7**. From these examples, we found that G4s formed in many common loci regardless of cell lines (**Figure S5**) and in some loci in a cell line-dependent manner (**Figure S5**, arrowheads), which indicated that the G4P detected the differences in G4 formation in different cell lines.

Among the 123,274 G4P peaks, 43,752 had a fold change ≥5 in signal relative to input (**Figure 5D**). The number of peaks in the four genomic regions dropped rapidly with an increase in the fold change of the peaks. Those peaks with high fold change should represent more stable G4s that form in higher frequency and/or last longer once formed. When the peaks in each genomic region were plotted as a percent of the total, those in the TSS±2kb region rapidly increased while those in the other three regions decreased (**Figure 5E**). This unique feature implied a more robust G4 formation in promoters driven by transcription. An obvious increment started at 4-folds, suggesting efficient G4 formation at this enrichment occurred in response to transcription. Taking ≥5 fold as a threshold, ~80% of the G4s were associated with genes (**Figure 5E**, gene±2kb, black curve). Owing to the concentration of PQS at promoters, G4s were detected in a much greater fraction in the TSS±2kb region than in the other three ones (**Figure 5F**). Counting peaks of ≥5 fold change, G4s were detected in >60% promoters (TSS±2kb) and ~70% genes. A summary **Table S4** provides details on the G4P peaks regarding their genomic coordinates, fold change, PQS motifs covered, and genes associated. Taken together, our data revealed that G4s readily form in association with transcription in living animal cells and promoters are their main playgrounds.

### Co-localization of G4P with native G4-binding proteins

The identification of G4P binding peaks allowed us to compare the binding activity of the G4P among the human cell lines and with several endogenous native proteins, namely FUS, TAF15, RBM14, and TARDBP, that have been reported to bind DNA and RNA G4s in vitro (33–35). Thus, we plotted distribution profiles of the G4P and the four G4-binding proteins around the G4P peak regions from the A549 cells (**Figure 6A**, top panels). We can see that the G4P-binding signal was also present in the other three human cell lines although their magnitudes varied (**Figure 6A**, top panels 1-4). In addition, binding signals were also found for the four native G4-binding proteins in these G4P-binding regions (**Figure 6A**, top panels 5-8), suggesting they might co-localized with the G4P at their binding sites. The formation of G4s in one DNA strand leaves its complementary cytosine-rich (C-rich) partner single-stranded. In correlation with this, the binding signal was also seen for two such proteins, i.e. PCBP1 and hnRNP K (**Figure 6A**, top panels 9-10) that recognize single-stranded C-rich DNA (35, 36).

**Figure 6.**
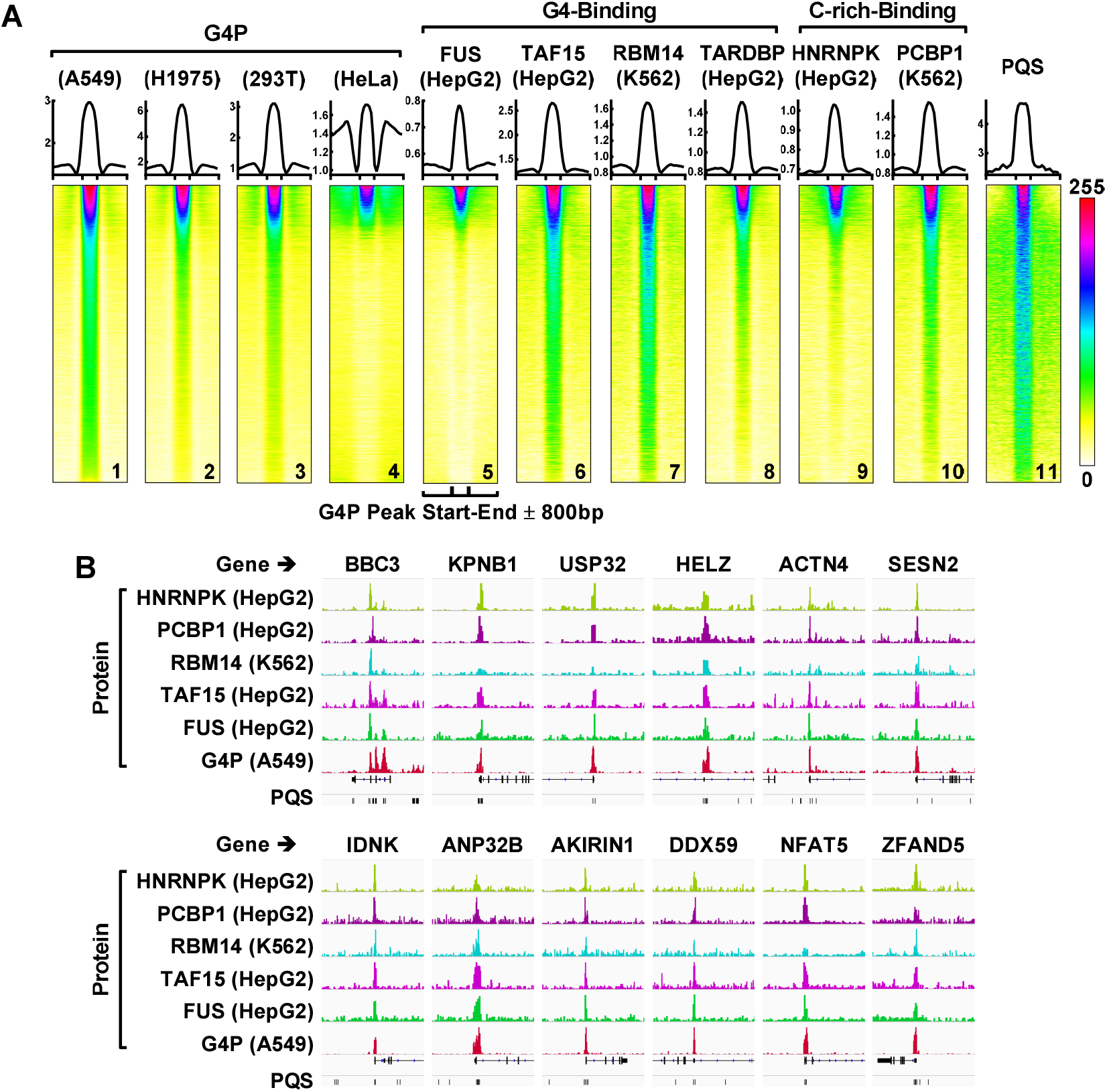
G4P binds genomic G4s in human cells as the indicated native G4-binding proteins do. (A) Profiles (top) and heatmaps (bottom) of proteins at the G4P peak ± 800 bp regions. The cell line is indicated in parenthesis. All heatmaps were produced over the G4P peak bed file of the A549 cells sorted on the mean G4P reads of the A549 cells in descending order. Values in each heatmap were normalized independently to the range of 0-255 to enhance the visual resolution. The bigwig files of the G4- and C-rich-binding proteins were from the Encode database (https://www.encodeproject.org/) with the following identifier numbers: ENCFF274FNN, ENCFF061RJA, ENCFF707TKH, ENCFF470CIC, ENCFF292JAR, and ENCFF882OWS. (B) Examples of protein binding peaks at G4P regions.

To obtain more insight, we sorted the G4P peak region file of the A549 cell line on the mean G4P reads in descending order and plotted heatmaps of the three groups of proteins over this G4P-binding region file. The results turned out that all the binding activities generally followed a similar gradient (**Figure 6A**, bottom panels 1-10) in a positive correlation with the presence of PQS motifs (**Figure 6A**, bottom panel 11). Clearly, these results showed that the G4P bound G4s as the native proteins did in cells, for which representative examples are shown in **Figure 6B**. They also revealed that a common landscape of G4 formation was largely shared among the different cell lines (**Figure 6A**, bottom panels 1-4) although their magnitudes varied (**Figure 6A**, top panels 1-4).

### Effect of G4P expression on transcription

Owing to the involvement of G4s in transcription, the expression of an exogenous G4-binding G4P in cells affects gene expression. In return, the changes in gene expression brought by the G4P, if any, are expected to affect G4 formation in genes because of its dependence on transcription activity (**Figure 3H**). To evaluate the impact of G4P, we assessed the changes in gene expression by RNA-Seq (**Figure S8**), revealing bi-directional changes in RNA level among genes. As shown in **Figure S8C**, the changes were no more than 2 folds in ~95% of the genes in the 293T cells with the G4P knock-in. For the plasmid-transfected cells, the RNA levels changed by less than 4 folds in >85% of the genes (**Figure S8**), slightly greater than that in the 293T cells, which might be explained by a greater expression with the transfection.

Gene expression is dynamic and exhibits intrinsic temporal fluctuations of up to >10-fold changes (36,37), which significantly overweighs the changes in gene expression brought by the G4P (**Figure S8**). This suggested the introduction of G4P might not have an overwhelming impact on the global gene expression of the cells. On the other hand, there are dozens of native G4-binding activities in cells (38); some can bind G4s more tightly than G4P with a *K*_d_ at nM (33) or even pM (39) level. The expression of the G4P was merely adding one more binding activity to the many ones that already existed, such that the G4P had to compete with them to bind G4s. Nevertheless, the fact that G4P binding largely co-localized with the targets of the native binding proteins in the cells that did not express G4P (**Figure 6**) suggested the G4P-G4 interaction generally reflected the landscape of G4 formation in the cells. For these reasons, we assume that introducing G4P might affect the magnitude of G4 formation to a limited extent in most of its targets unless a gene was completely switched on or off by the G4P. The general conclusion regarding G4 formation should remain qualitatively valid.

### A brief comparison with the BG4 antibody

To briefly compare with the G4 detection in fixed cells with the popular BG4 antibody (3), we looked at the G4P reads distribution around TSS and the PQS coverage in the G4P peaks in all the cells we tested. The results revealed differences in three general aspects in comparison with the BG4. First, the profiles of G4P reads all showed two peaks (**Figure 7A**) in correlation with the two peaks of PQSs (**Figure 3G**, insert) and the asymmetric transcription activity at TSS while those of the BG4 all appeared as a single symmetric peak centered at TSS (**Figure 7B**). Second, the enrichment peak at the TSS of the G4P was generally much higher than that of the BG4. Third, the percentage of PQS-positive peaks detected by G4P (**Figure 7C**) was significantly greater than those detected by the BG4 (**Figure 7D**). These differences demonstrated significant improvements in resolution, sensitivity, specificity, and efficiency of the G4P in recognizing G4s, which might be attributed to G4P’s tiny size and high affinity to G4s.

**Figure 7.**
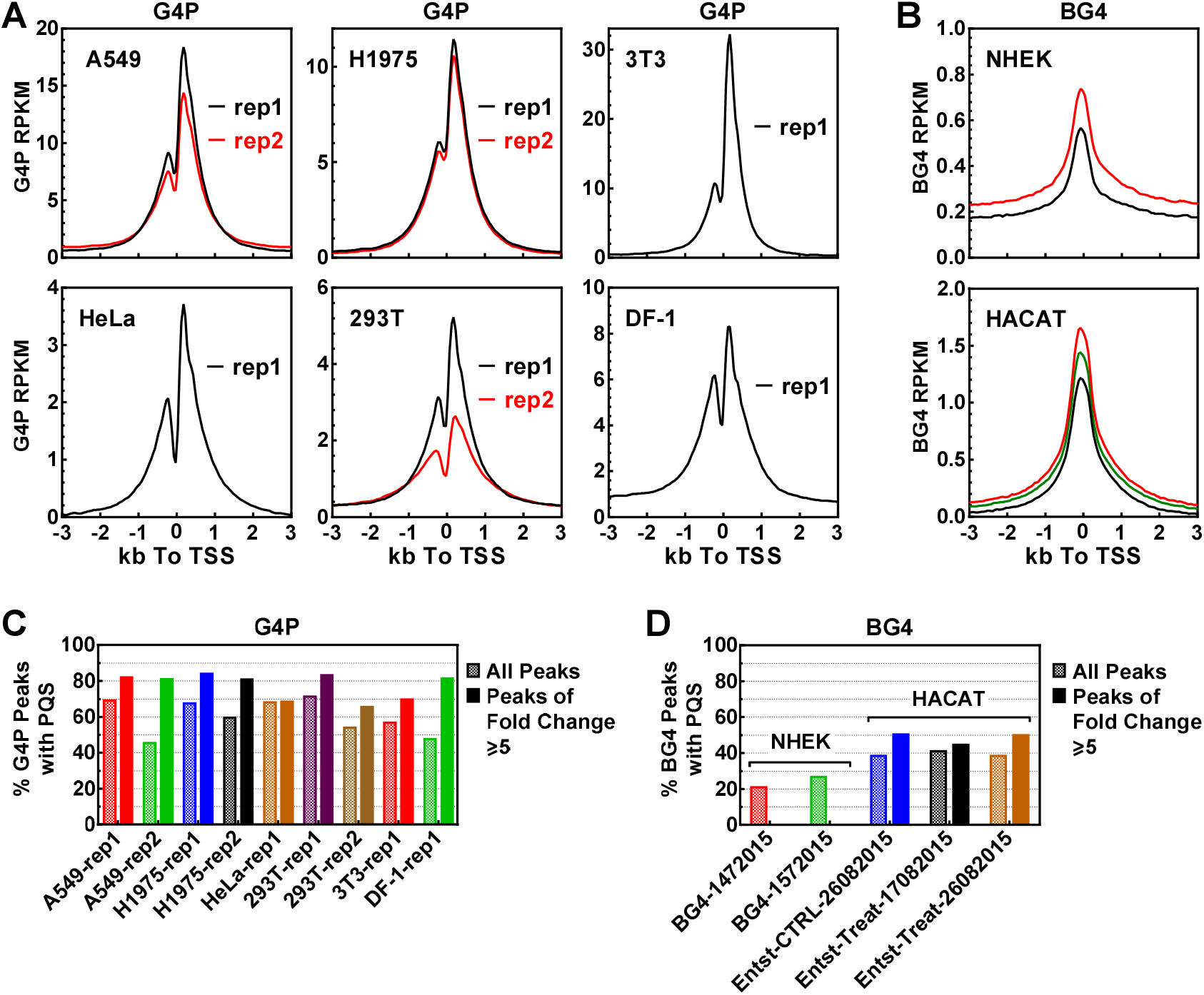
Comparison between detection of G4s in living cells using G4P and in fixed cells using BG4. Distribution of (A) G4P or (B) BG4 reads around TSSs. Percent of (C) G4P or (D) BG4 peaks that overlapped with at least one PQS by one or more nucleotides. Fastq and narrow peak files for BG4 were downloaded from GSE76688 in the GEO database (https://www.ncbi.nlm.nih.gov/geo/). Bigwig files were produced from the fastq files as were the files for G4P using the same parameters whenever applicable. Calculation of distribution and PQS overlapping were all conducted in the same way. The cell line is indicated under the X-axis or in the panel.

## Discussion

In summary, we developed a unique probe to capture G4 formation in living cells with the ability to reveal the locations, magnitudes, and sequence identities of G4s in a whole genome (**Table S4**). Using this probe, we observed a robust formation of G4s in genes as a native event in transcription in animal cells (**Figure 3–6**), which establishes a molecular basis for the role of G4s in transcription. The positive correlation of G4 formation with transcription activity (**Figure 3H**) and PQS load (**Figure 4**, A and E) confers a capability for the G4-related regulation to be performed in response to the activity of transcription defined by the amount and type of PQS motifs. The similar landscapes of G4 formation shared among the different cell lines (**Figure 5A**, **Figure S3**, and **Figure 6**) suggest the presence of a common and essential mechanism involving G4s in transcription regulation.

As a G4 probe, the G4P possesses several key traits that outperform antibodies and native G4-binding proteins. In comparison, the G4P has a much smaller size, higher specificity, and affinity (*K*_d_ of low-nM). Most importantly, it overcomes the problem of impermeability and disulfide bonds associated with antibodies such that the G4P is readily applicable in living cells. With the removal of >90% of the amino acid residues from the original RHAU, the G4P is unlikely to interact with other proteins as the RHAU and other proteins might do, therefore, ensuring direct target recognition and specificity. While native G4-binding proteins can also locate G4 formation, they may result in false-positive detection because of potential interaction with other macromolecules. For instance, the hnRNP A2* protein can bind the RNA component of telomerase in addition to telomere DNA G4s (40). In theory, it may crosslink genomic DNA at where the RNA component of telomerase is synthesized when a cell is fixed. On the other hand, the tiny size of the G4P helps it gains higher resolution and accessibility to G4s. The core G4P is only 6.7 kDa (spanning only two RHAU23s), being about half the size of a nanobody (12-15 kDa, the smallest antibody derivative). Even with the NLS and FLAG tags, it is merely 11 kDa, still smaller than a nanobody, not to mention the scFv (27 kDa) and regular antibody (~150 kDa). Chemical probe has been used for detecting non-B DNA structures in living cells by ChIP-Seq (41) that covers a broader range of secondary structures including G4s. Our G4P is dedicated to G4s and expected to provide unique complementation to such existing tools.

The G4P also offers many opportunities to expand its functionality in various applications both in *vitro* and in vivo to study or modify the activity of G4s and related biological and medical processes. For instance, the fusion of G4P with fluorescent proteins may enable visualization of G4s and G4-related activities. Fusion with enzymes may enable modifications to protein/DNA/RNA or create desired reactions in association with G4s that would not occur natively. G4P-ChIP may be employed to identify proteins or processes that interact or associate with G4s. The 64-aa core G4P may be used as building blocks and find application in biological drugs, molecular nanodevices to confer G4 recognition capability. The core G4P can be synthesized in large quantities at low cost with a possibility of chemical modifications during or after synthesis. Functional groups or labels, such as radionuclei, quantum dots, nanoparticles, fluorescent dyes, etc., can be conjugated through the amine or carboxyl groups of the G4P. Collectively, all these beneficial attributes make the G4P a satisfactory G4 affinity agent.

## ACKNOWLEDGMENTS

This work was supported by the National Natural Science Foundation of China, grant numbers 21672212, 21432008, 21708042, and 21602220.

## Supporting Information

### SI Materials and Methods

#### Identification of PQS motifs in genomes

Chromosome files (hg19 or hg38 for human, mm9 for mouse, and gal5 for chicken) in fasta format were downloaded from the UCSC website (http://genome.ucsc.edu/). Global searching of four types of PQS motifs (i.e. 4G, 4GL15, Bulge, and GVBQ) was performed using home-made Perl scripts. 4G PQS motifs were identified using a regular expression G{3,}(.{1,7}?G{3,}){3,} as previously described (1,2). Identification of the other three PQSs involved a first round of search to find motifs with the desired feature. Each motif found was then subjected to several additional rounds of pattern matching to remove unwanted motifs. A flow chart and the variables for the identification of 4GL15 PQS is shown below as an example.

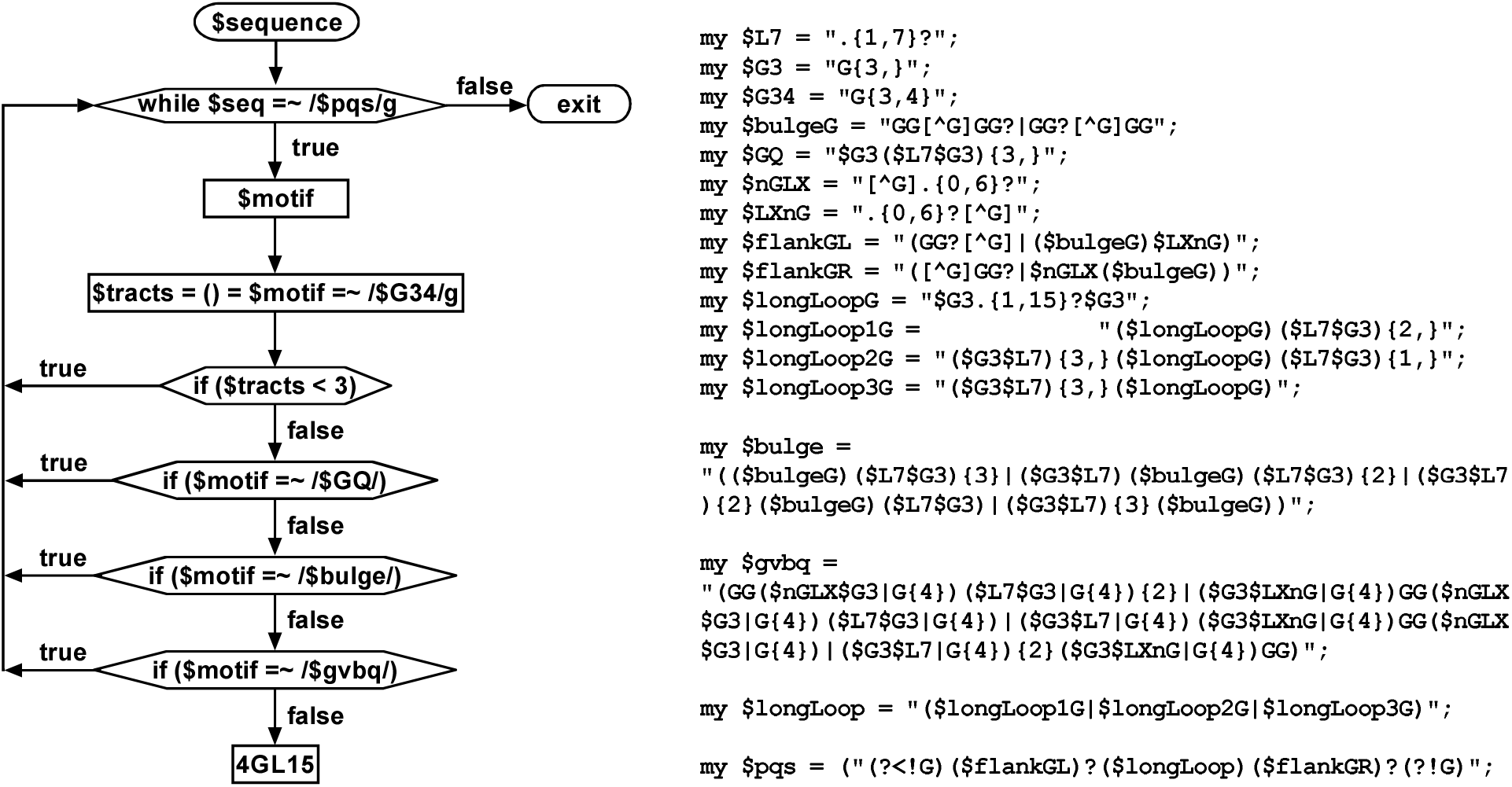

#### Plasmid construction for G4P-ChIP

DNA coding the RHAU23-(GTGSGA)-3xFLAG (RHAU23) or RHAU23-(GTGSGA)3-RHAU23-(GTGSGA)-3xFLAG were synthesized by Generay Biotechnology (Shanghai, China) and inserted into pET28b between the Nde I and EcoR I sites to obtain pET28b-RHAU23 and pET28b-G4P. The former coding sequence was also inserted into pIRES2-EGFP between the Nhe I and EcoR I sites to obtain pG4P-IRES2-EGFP. The DNA fragment containing a nuclear localization signal (NLS) of the SV40 large antigen (PKKKRKV) was synthesized by Sangon (Shanghai, China) and inserted into pG4P-IRES2-EGFP at the Nhe I site to obtain plasmid pNLS-G4P-IRES2-EGFP. The construction of plasmids for stable transfection in cell lines was as described (3). The DNA fragment expressing NLS-G4P and eGFP was amplified from the pNLS-G4P-IRES2-EGFP and inserted into the AAVS1 donor plasmid between the Spe I and Sal I sites. The AAVS1 loci specific guide RNA sequence (5’-GTCACCAATCCTGTCCCTAG-3’) was designed by the online CRISPR tool (crispr.mit.edu.) and inserted into the PX330 plasmid between two Bbs I sites (4,5).

#### Cell lines and cell culture conditions

A549, NCI-H1975, 293T, HeLa-S3, 3T3, and DF-1 cells were kindly provided by Stem Cell Bank, Chinese Academy of Sciences. Cells were grown in DMEM supplemented with 10% FBS and 1× penicillin-streptomycin.

#### Transient transfection and gene knock-in

For transient transfection, cells were cultured in 15 cm dishes to 70-80% confluence and transfected with 30 μg pNLS-G4P-IRES2-EGFP using lipofectamine 3000 (Thermo Scientific) according to the manufacturer’s instructions. Cells were cultured for an additional 24 hrs before harvesting. For stable transfection, AAVS1 donor and PX330 plasmid containing G4P and AAVS1 gRNA were co-transfected into 293T cells using lipofectamine 2000 (Thermo Scientific). After 24 hours, GFP positive single cell was sorted by a flow cytometer (MoFlo XDP, Beckman) into a 96-well plate. Cells were cultured for two weeks and the cell lines with stable expression of G4P were verified by PCR, western blot, and immunofluorescence.

#### Recombinant G4P and RHAU23

The plasmid pET28b-G4P and pET28b-RHAU23 were transformed into the E.coli strain BL21 (DE3). Cells were grown in LB medium supplemented with 0.05 mg/mL kanamycin at 37 °C for 3 hrs. G4P and RHAU23 expression were induced with 1 mM isopropyl thiogalactoside for 4 hrs at 37 °C. The two peptides were purified using the Capturem HIS-tagged Purification Miniprep Kit (Takara, Dalian) and diluted in buffer containing 20 mM Tris-HCl (pH 7.4), 150 mM NaCl, 0.1 mM EDTA, 50% glycerol, and stored at −20 °C.

#### Circular dichroism (CD) spectroscopy

CD spectroscopy was conducted as described (6). The incubation of DNAs (**Table S1** and **S3**) with G4P was carried out at RT or the indicated temperature and time interval with a 1:1 protein/DNA ratio.

#### Electrophoretic mobility shift assay (EMSA)

DNAs (**Table S1** and **S3**) were dissolved at 10 nM in a buffer containing 20 mM Tris-HCl (pH 7.4), 75 mM KCl, 1 mM EDTA,0.4 mg/mL BSA, without or with (in case of dsDNA) 40% (w/v) PEG 200, denatured at 95 °C for 5 min, and slowly cooled down to 25 °C. DNA was then incubated with G4P of the indicating concentration at 4 °C for 1 hour. Samples were resolved on 12% non-denaturing polyacrylamide gel containing 75 mM KCl in the absence or presence (in case of dsDNA) of 40% (w/v) PEG200 at 4 °C for 2 hours in 1×TBE buffer containing 75 mM KCl. DNA was visualized by the FAM dye covalently labeled at their 5’ end of the DNA on a ChemiDoc MP (Bio-Rad) and digitized using the Image Quant 5.2 software. Dissociation constant was determined by fitting the fractional bound DNA (Y) in a 1:1 stoichiometry against the free G4P concentration (X) to the equation Y=F1×X/(*K*_d1_+X)+(1-F1)×X/(*K*_d2_+X), where F1 denoted the fraction of the subpopulation associated with the *K*_d1_.

#### Pull down of G4-containing plasmid with G4P

Plasmids containing the indicated PQS or a mutant control sequence (**Table S2**) on the non-template strand were constructed as described (7). Transcription was carried out in a total volume of 50 μl at 37 °C for 1 hour in transcription buffer containing 40 mM Tris-HCl (pH 7.9 at 25 °C), 8 mM MgCl_2_, 10 mM DTT, 2 mM spermidine, 50 mM KCl, 200 U T7 RNA Polymerase (Thermo Scientific), 40 U RNase inhibitor (Thermo Scientific), 2 mM NTP, 0.5 μg pEGFP-N1 plasmid, and 0.5 μg PQS or control plasmid. The reaction was stopped by addition of EDTA to 16 mM. 1 μl of samples were used as input and diluted in 50 μl water. The remaining samples were mixed with 0.2 μM G4P and incubated at 4 °C for 30 min.

For the pull-down assay, 12 μl anti-FLAG M2 magnetic beads (Sigma-Aldrich) were washed three times with binding buffer (20 mM Tris-HCl, pH 7.4, 150 mM KCl, 1 mM EDTA, and 0.5% Triton X-100) and then resuspended in 200 μl binding buffer. The beads were then incubated with the DNA samples on a rotating mixer at 4°C for 2 hours. After the removal of the supernatant, the beads were washed ten times with the binding buffer accompanied with three times of transfers to new tubes. The precipitated plasmids were released from the beads by incubating with 0.3 mg/ml 3xFLAG peptide (Sigma-Aldrich) in a 50 μl binding buffer at 4°C for 0.5 hours. The released plasmid and the input samples were treated with RNase A at 37°C for 15 min and proteinase K at 60°C for 20 min, followed by inactivation at 95°C for 10 min.

qPCR was conducted on the CFX Connect thermocycler (Bio-Rad) for the PQS or control plasmid using a PCR primer pair of 5’-TCCGACACTATGCCATCCTGA-3’ and 5’-TGCAGTCGTTTCGTATCGTTGA-3’. The EGFP gene on the pEGFP-N1 plasmid using a primer pair of 5’-GCAGAAGAACGGCATCAAGG-3’ and 5’-CGGACTGGGTGCTCAGGTAG-3’. After calibrating with the internal control, the enrichment was calculated using input DNA as a reference.

#### G4P-ChIP library construction

Approximately 0.5-1 ×10^7^ transiently or stably transfected cells expressing G4P were crosslinked with 1% formaldehyde for 20 min at room temperature. Fixation was quenched by 0.125 M glycine for 15 min. The fixed cells were washed twice with PBS, suspended in NP-40 buffer (10 mM Tris-HCl pH 7.4, 150 mM NaCl, 0.5% NP-40 and 2 mM AEBSF) and incubated on ice for 10 min. After centrifugation at 800×g for 5 min, the cell pellet was resuspended in a CHAPS buffer (20 mM Tris-HCl pH 7.4, 0.5 mM EGTA, 50 mM NaCl, 0.5% CHAPS, 10% glycerol and 2 mM AEBSF). The suspension was incubated on ice for 30 min and centrifuged at 800×g for 5 min. The pellet was resuspended in 1 ml 1× dsDNase digestion buffer supplied with 50 μl dsDNase (Invitrogen, EN0771) and incubated at 37 °C for 20 min with constant agitation. A final concentration of 20 mM EDTA was added to terminate the reaction. The nuclei were pelleted by centrifugation at 15,000×g at 4°C and the supernatant was incubated on ice. The nuclei pellet was resuspended in 500 μl wash buffer (150 mM NaCl, 10 mM Tris-HCl, pH 7.4, 0.1 mM EDTA, 0.5% Triton X-100) and sonicated for 30-60 seconds in an ice-cold water bath. After centrifugation at 15000×g for 5 min, the supernatant was collected and combined with the supernatant from the previous step.

For library preparation, 50 μl of anti-FLAG M2 magnetic beads (Sigma-Aldrich) were washed with washing buffer (10 mM Tris-HCl, pH 8.0, 150 mM NaCl, and 0.5% Triton X-100) and blocked in the same buffer containing 75 μg/ml single-stranded sperm DNA and 1 mg/ml BSA. 1% chromatin fragment was saved as input and the remaining was incubated with blocked anti-FLAG magnetic beads in rotation at 4 °C for 3 hrs. Beads were sequentially washed ten times with washing buffer and transferred to new tubes three times. The chromatin was eluted with 300 μg/ml 3xFLAG peptide (Sigma-Aldrich) and incubated at 4 °C for 1 hr. The eluted chromatin and the input samples were incubated with proteinase K at 65 °C overnight. After sequential RNase A and proteinase K digestion, DNA fragment was cleaned by extraction with phenol:chloroform:isoamyl alcohol, followed by ethanol precipitation. Libraries were constructed from the recovered DNA fragment using the NEBNext Ultra II DNA LibraryPrep Kit from Illumina (NEB) according to the manufacturer’s instructions. The next-generation sequencing was performed with Illumina HiSeq X Ten by Genewiz (Suzhou, China).

#### RNA-Seq

Total RNA from cells expressing G4P was prepared using the SV Total RNA Isolation System (Promega). mRNA was purified from total RNA using the NEBNext Poly(A) mRNA Magnetic Isolation Module (NEB). RNA library was constructed using the NEBNext Ultra II Directional RNA Library Prep Kit from Illumina (NEB) according to the manufacturer’s instructions. The next-generation sequencing was performed with Illumina HiSeq X Ten by Genewiz (Suzhou, China).

#### ChIP-qPCR

Primers for amplifying PQS-proximal and distal regions were given in **Table S5**. qPCR reaction was performed using the GoTaq qPCR Master Mix (Promega) and qTOWER 2.2. The cycling condition was 95 °C for 20 s followed by 45 cycles of 30 s at 95 °C and 30 s at 60 °C. The enrichment of the genomic locus in the chip sample relative to the input was calculated using double delta Ct analysis with a PQS negative region as references.

---

**Figure S1.**
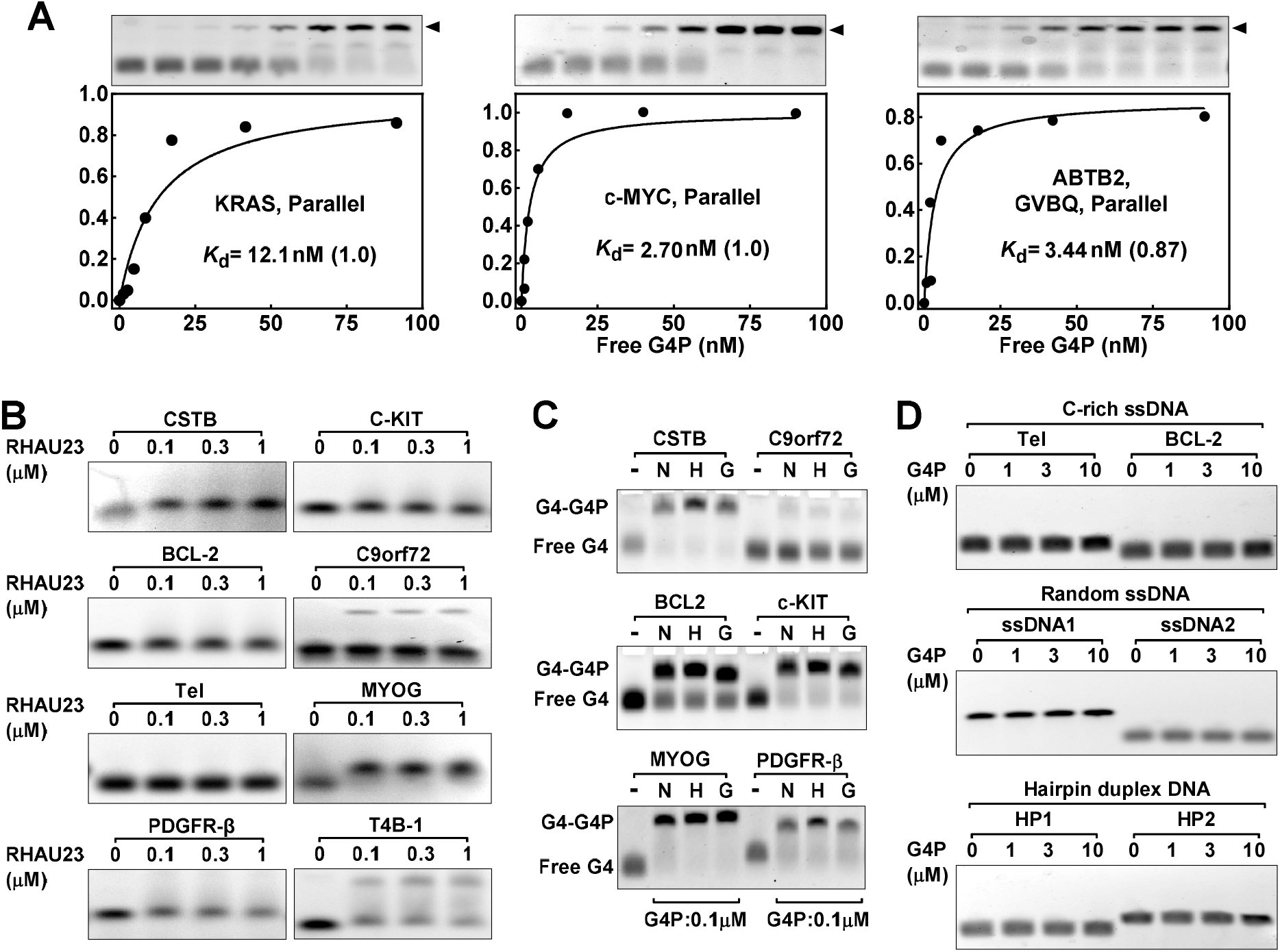
Additional characterization of RHAU23 and G4P by the electrophoretic mobility shift assay (EMSA). (A) Dissociation constant *K*_d_ of G4P to G4s. (B) RHAU23 bound poorly to G4s. (C) G4P binding to G4s with or without tags. G4P with tags was able to bind G4 as the G4P core. -: No G4P; N: G4P with an NLS-tag at the N terminal and a 3xFLAG tag at the C terminal; H: G4P with a HIS-tag at the N terminal and a 3xFLAG tag at the C terminal; G: G4P core without a tag. (D) G4P does not bind non-G4 DNA. Details of DNA can be found in **Table S1**.

**Figure S2.**
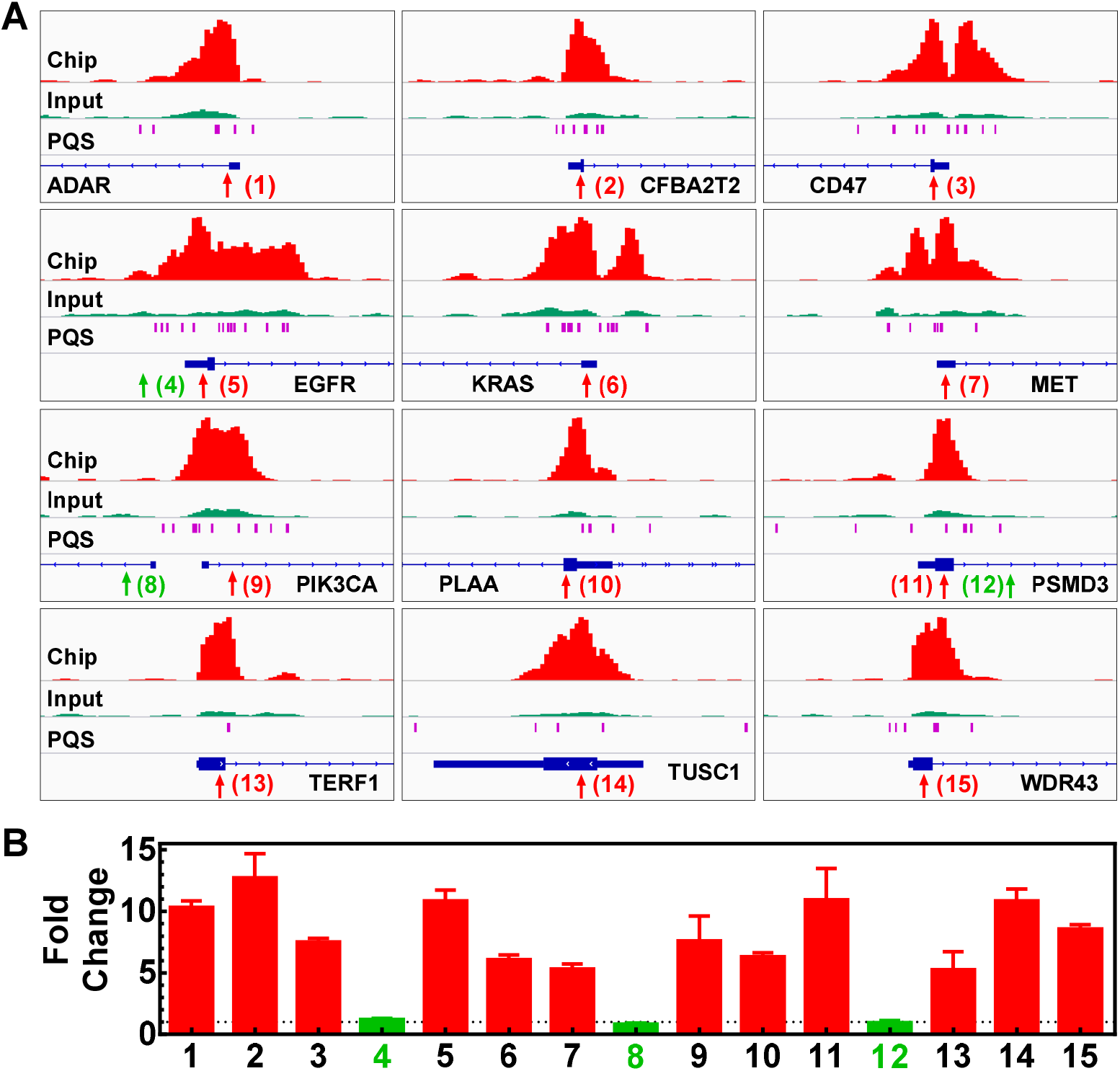
Verification of G4P enrichment in genomes of A549 cells by ChIP-qPCR. (A) qPCR regions are indicated by arrowheads. (B) Enrichment of G4P at indicated qPCR regions expressed as means of duplicate with range.

**Figure S3.**
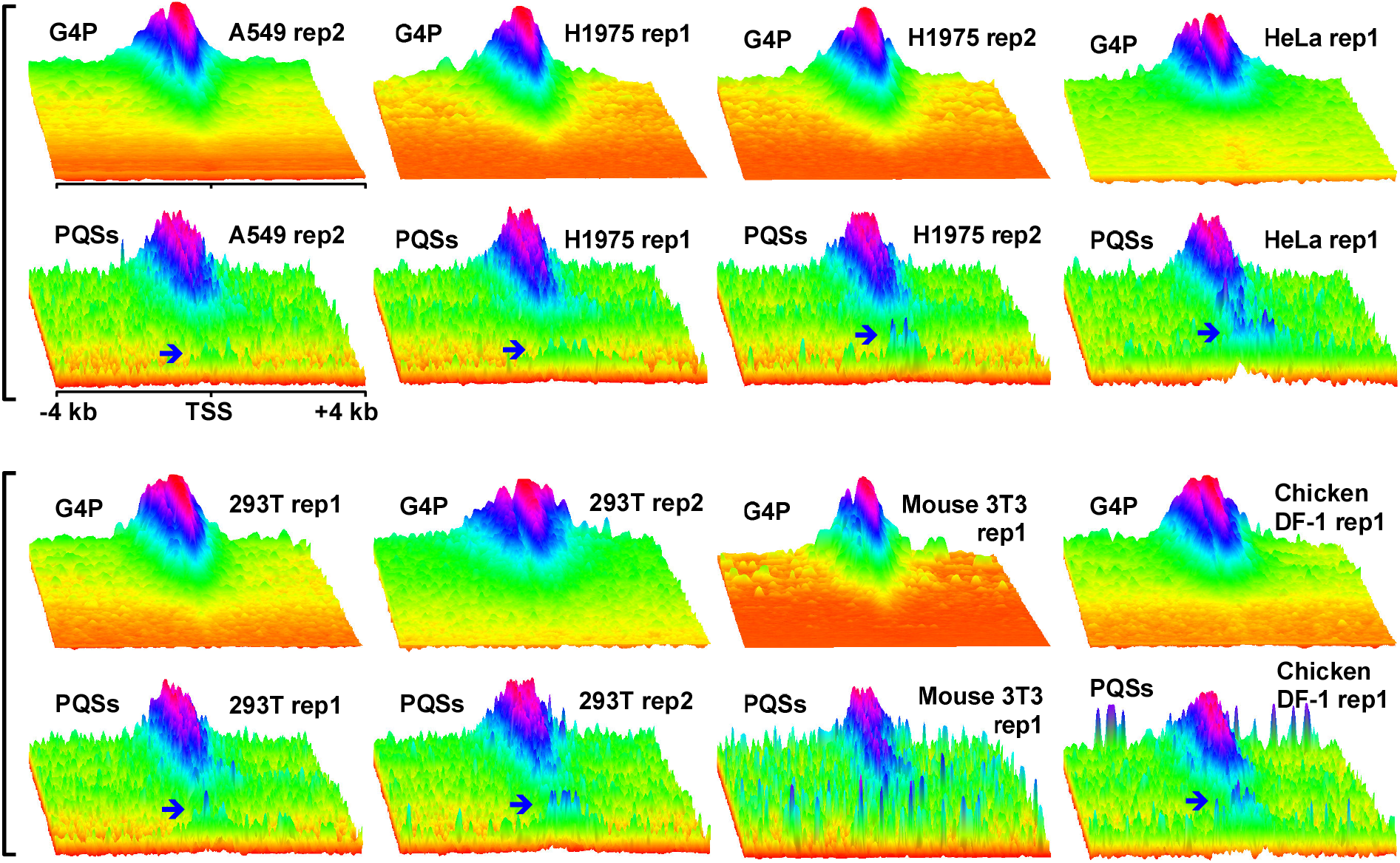
Overview of G4 formation and PQS distributions in A549, 293T, HeLa-S3, NCI-H1975, mouse 3T3, and chicken DF-1 cells in the TSS±4kb regions of RefSeq genes. Figure 5A and A549 rep2, 293T rep1, and rep2 were biological replicates (independent starting from cell culture); H1975 rep1 and rep2 were sequencing replicates. Data were analyzed as in Figure 5, A and B. The blue arrowhead shows a small fraction of the PQS motifs with a low probability of G4 formation.

**Figure S4.**
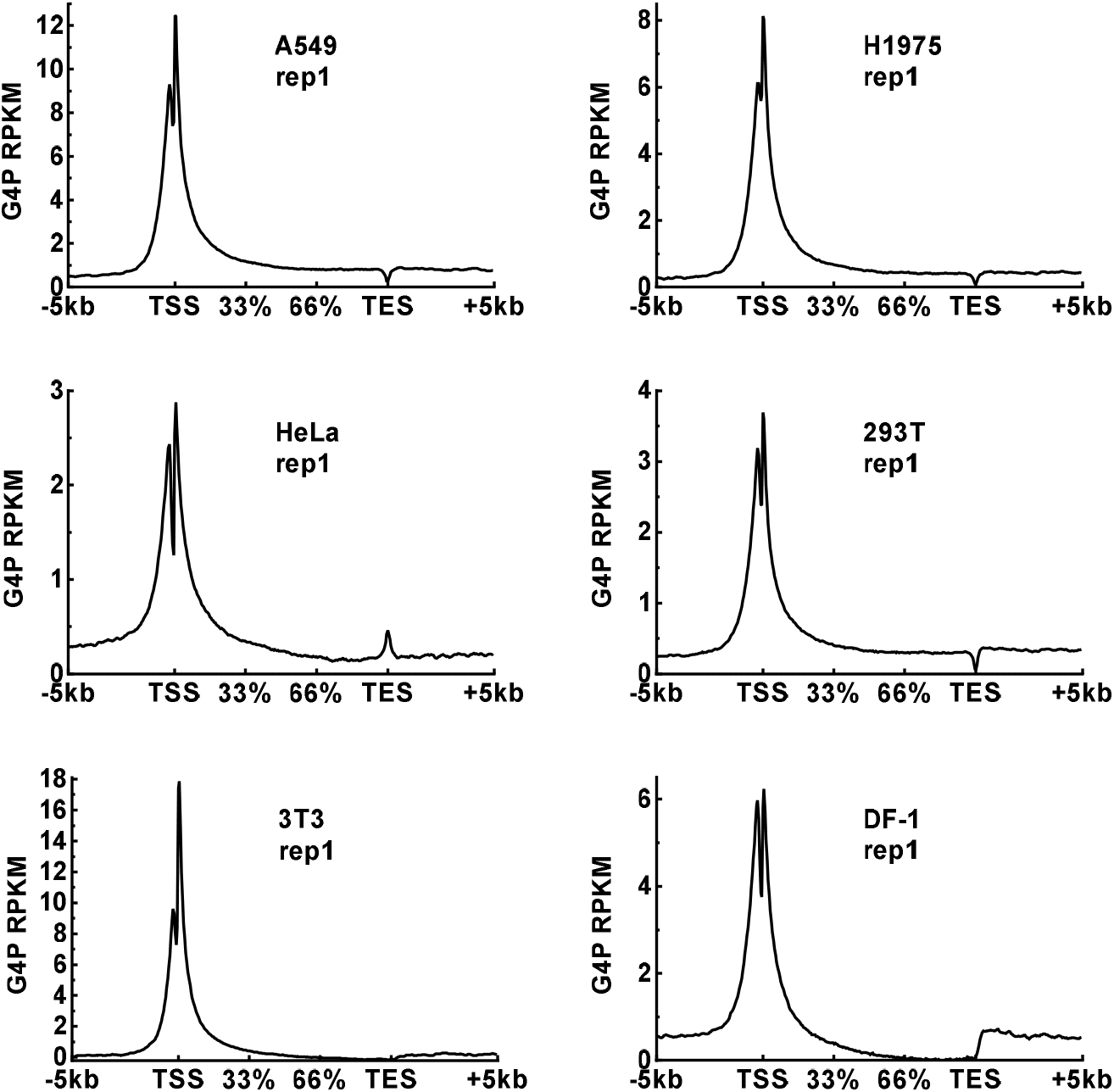
G4P reads distribution across RefSeq genes in A549, 293T, HeLa-S3, NCI-H1975, mouse 3T3, and chicken DF-1 cells.

**Figure S5.**
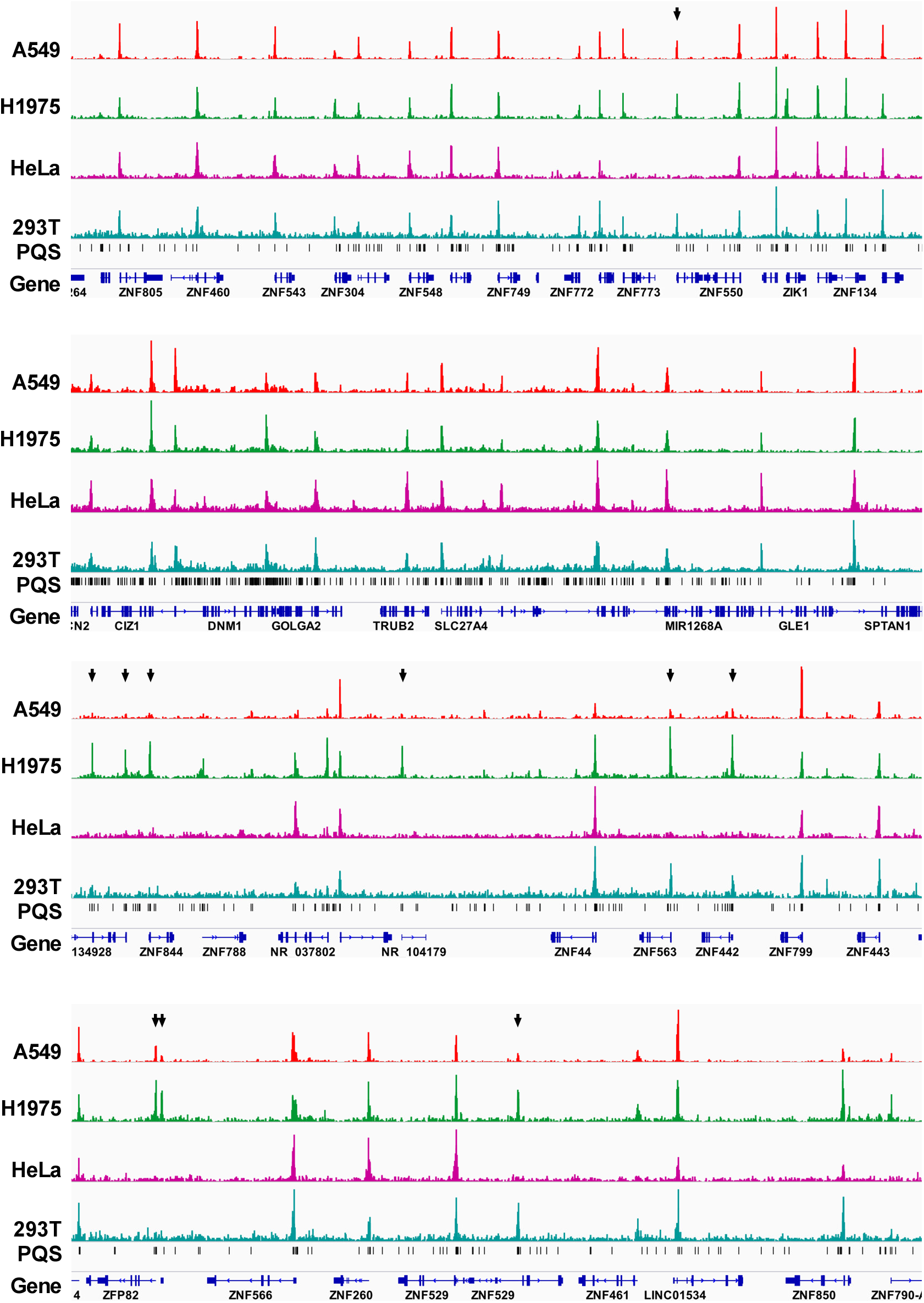
Examples of G4 formation in human cells depicted by the G4P peaks. Black arrowheads indicate loci where G4 formed in some cells but not in others.

**Figure S6.**
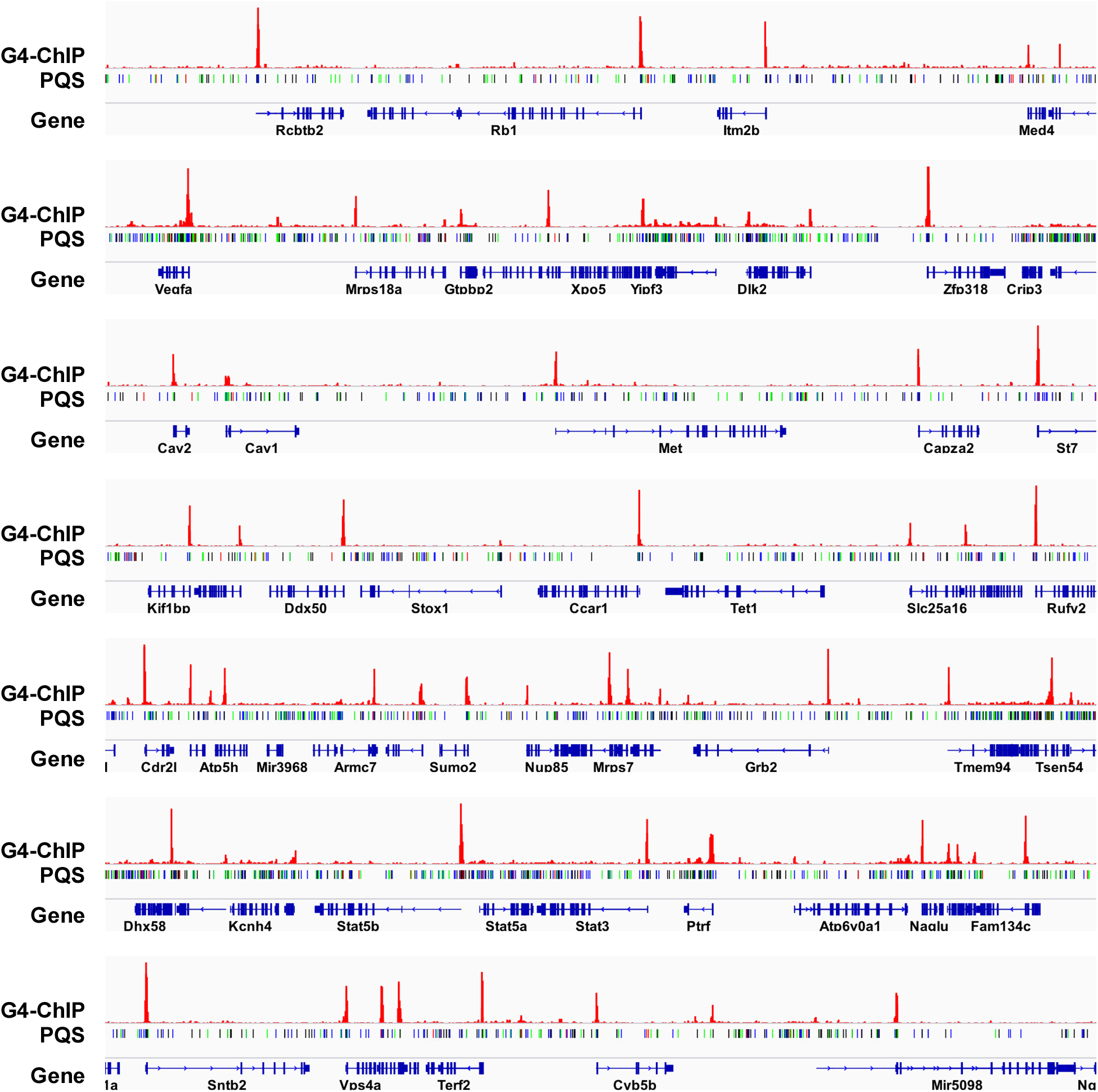
Examples of G4 formation in mouse 3T3 cells depicted by the G4P peaks.

**Figure S7.**
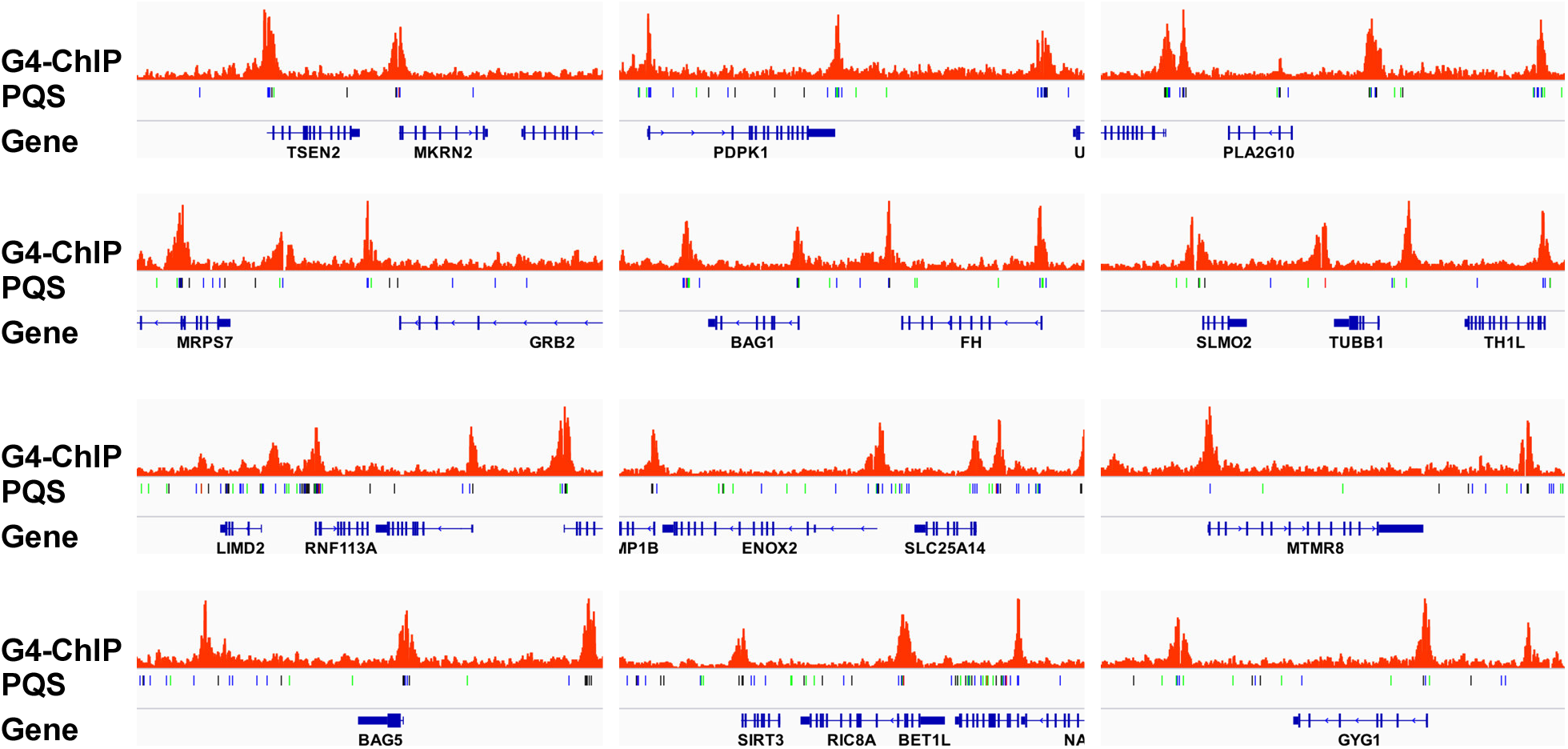
Examples of G4 formation in chicken DF-1 cells depicted by the G4P peaks.

**Figure S8.**
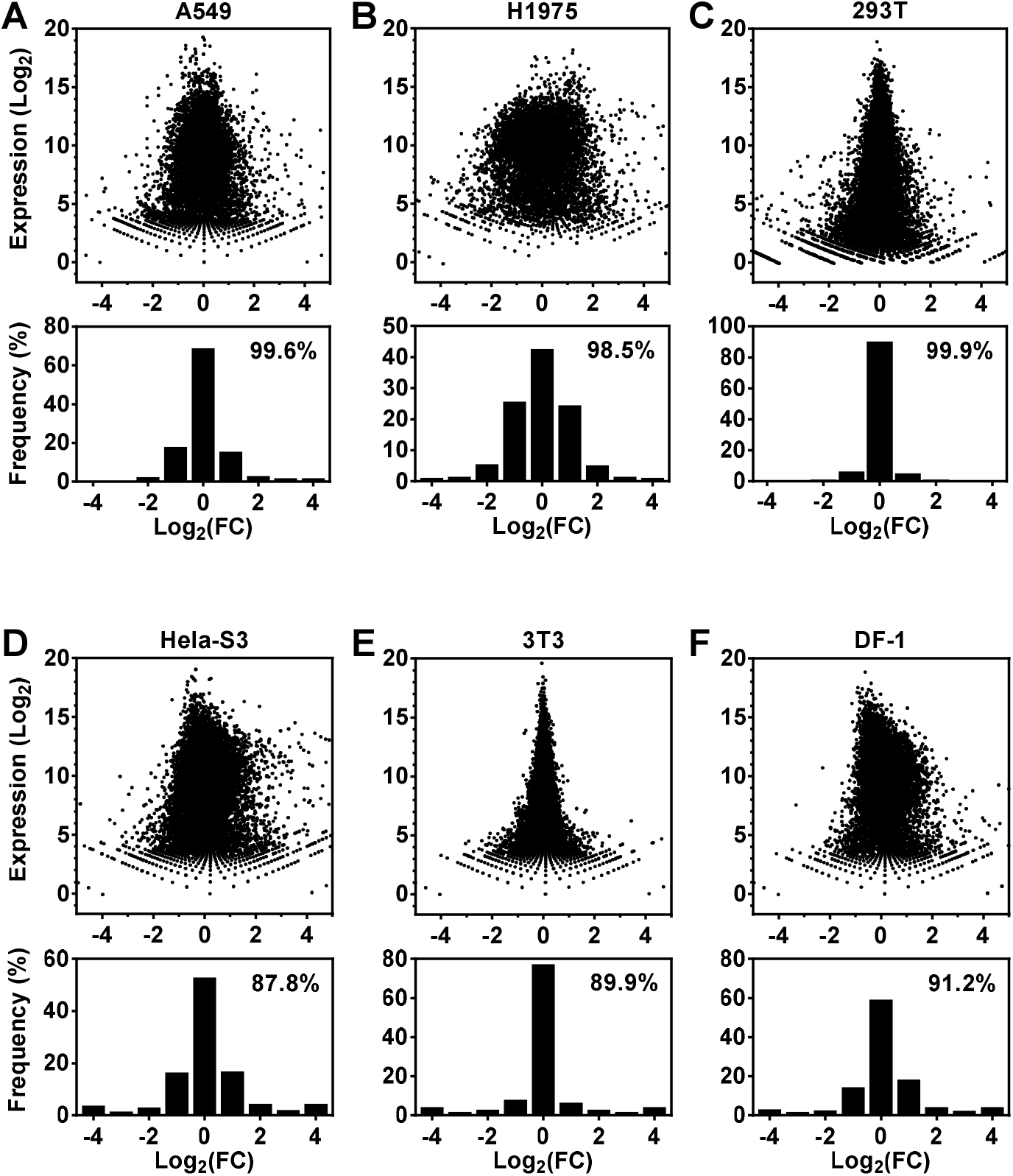
RNA-Seq results showing changes in gene expression in cells after the introduction of G4P. All cells, except the 293T, were transfected with a G4P expressing plasmid. G4P was introduced into 293T cells by sitespecific gene knock-in. Numbers inside the panels indicate the percentage of genes whose change in RNA level is less than or equal to 2^2^ (4 folds).

**Table S1.**
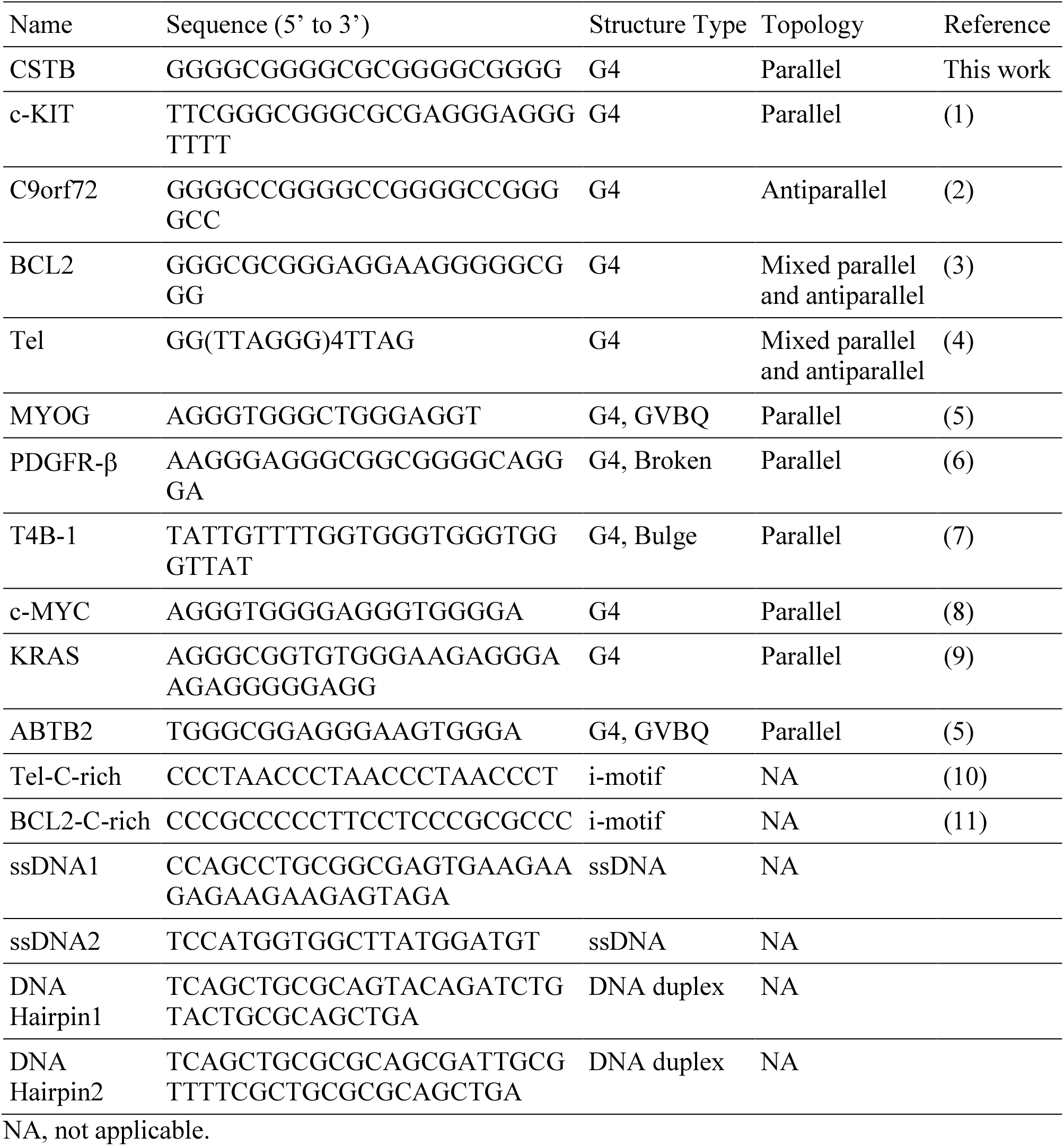
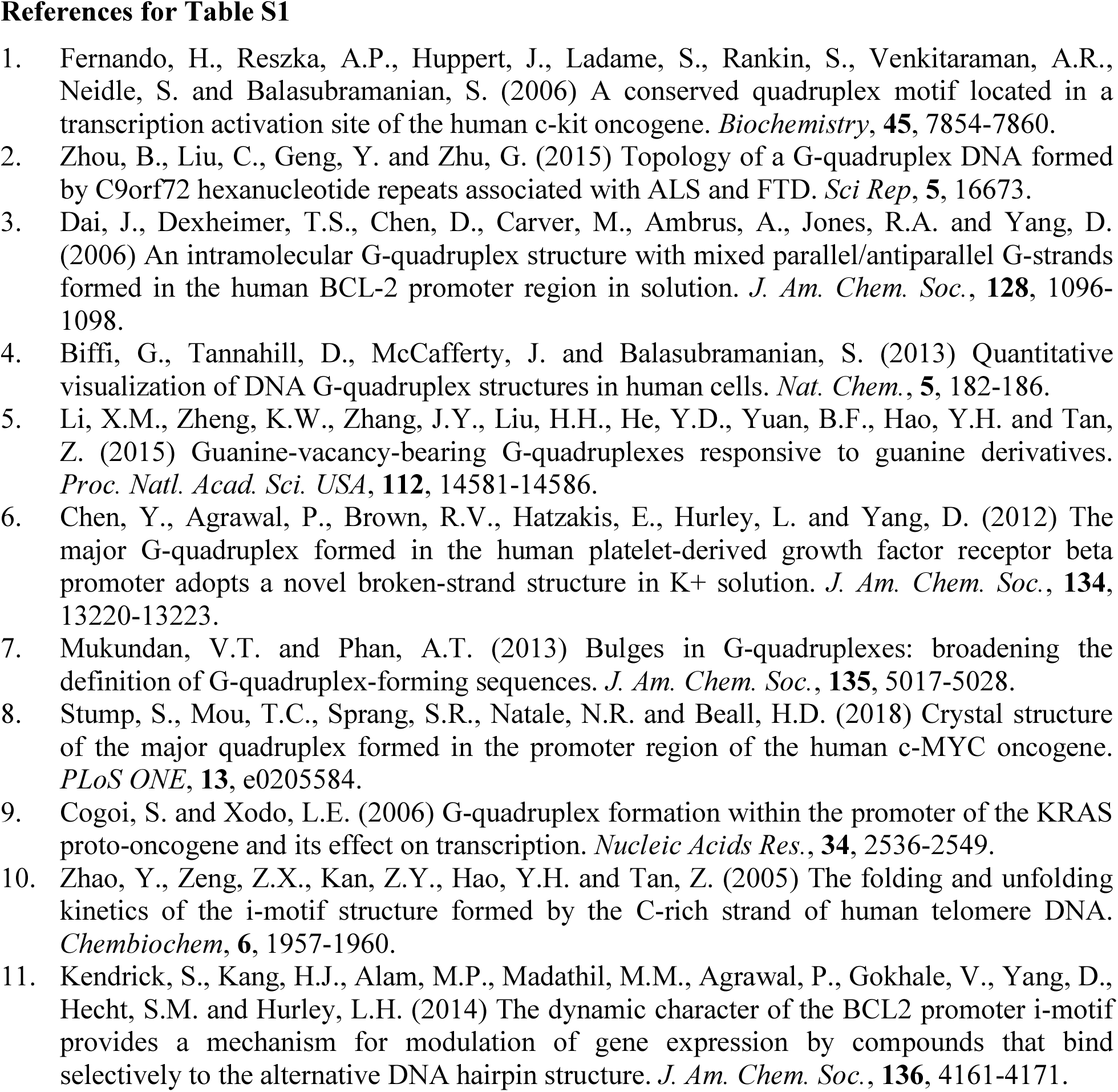
Sequences of DNAs used in binding and CD analysis.

**Table S2.**
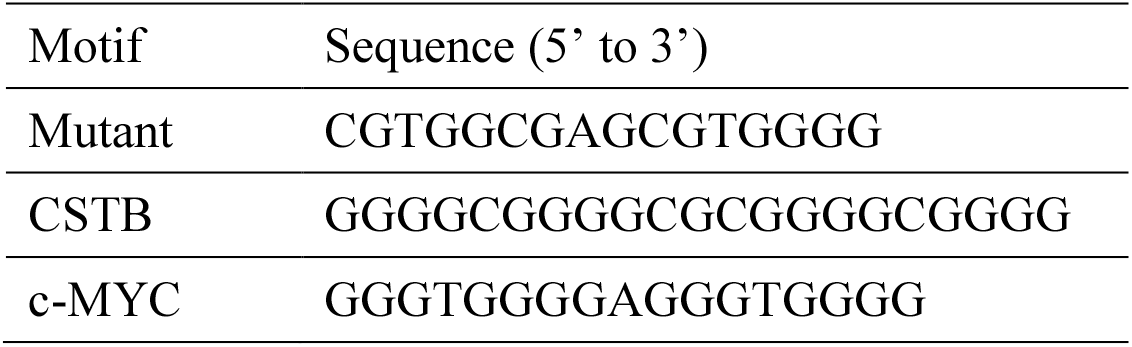
Sequences of PQS motifs used in plasmid pull-down.

**Table S3.**
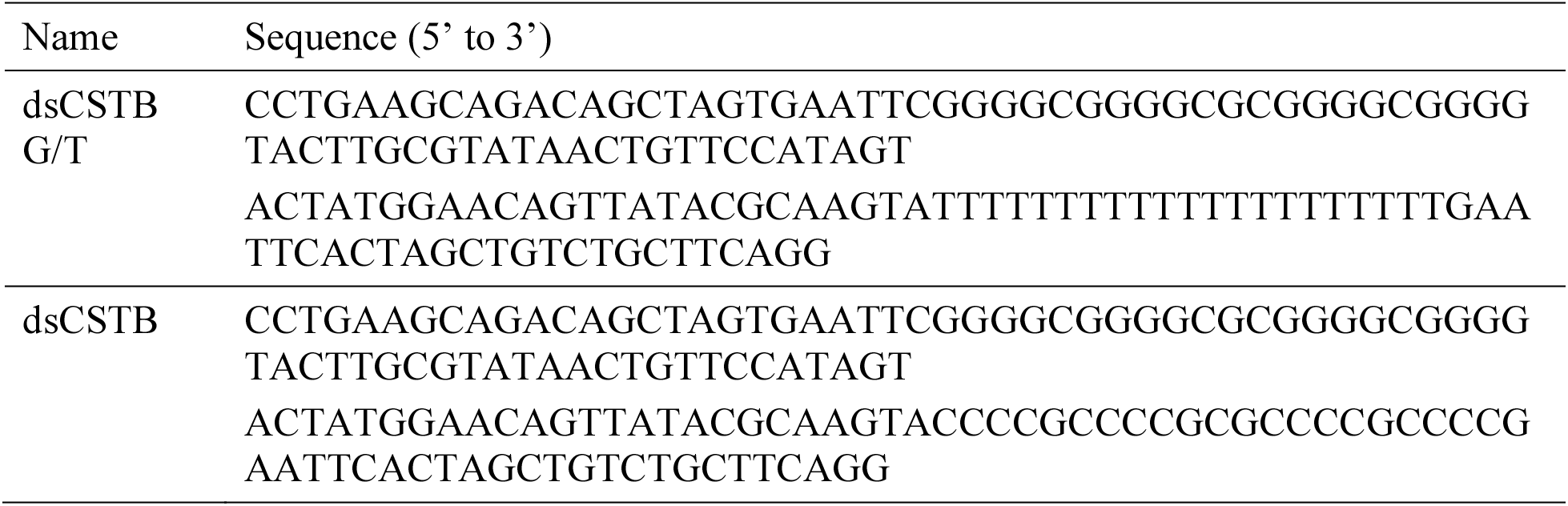
Sequences of dsDNA used in CD and EMSA analysis.

**Table S4.**
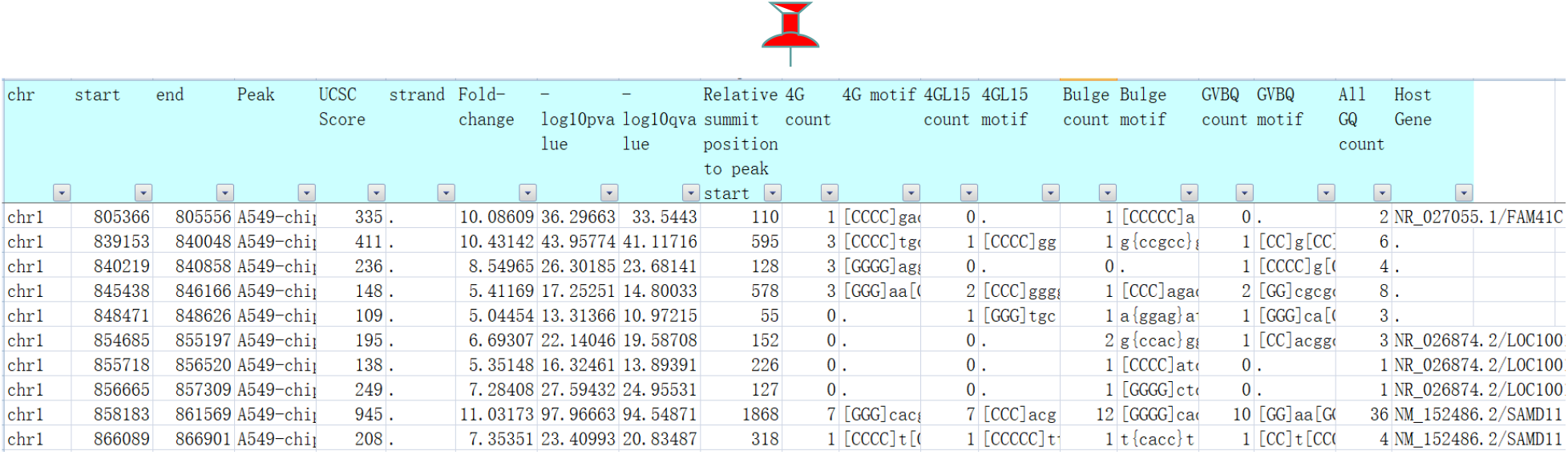
G4P peaks detected in human A549 cells (rep1). This table holds more than 1000 pages for the peaks of ≥5 fold changes, so it is provided in an embedded file: Table-S4-A549-rep1-G4P-narrowPeak-FCeg5-with-PQS.xlsb.

**Table S5.**
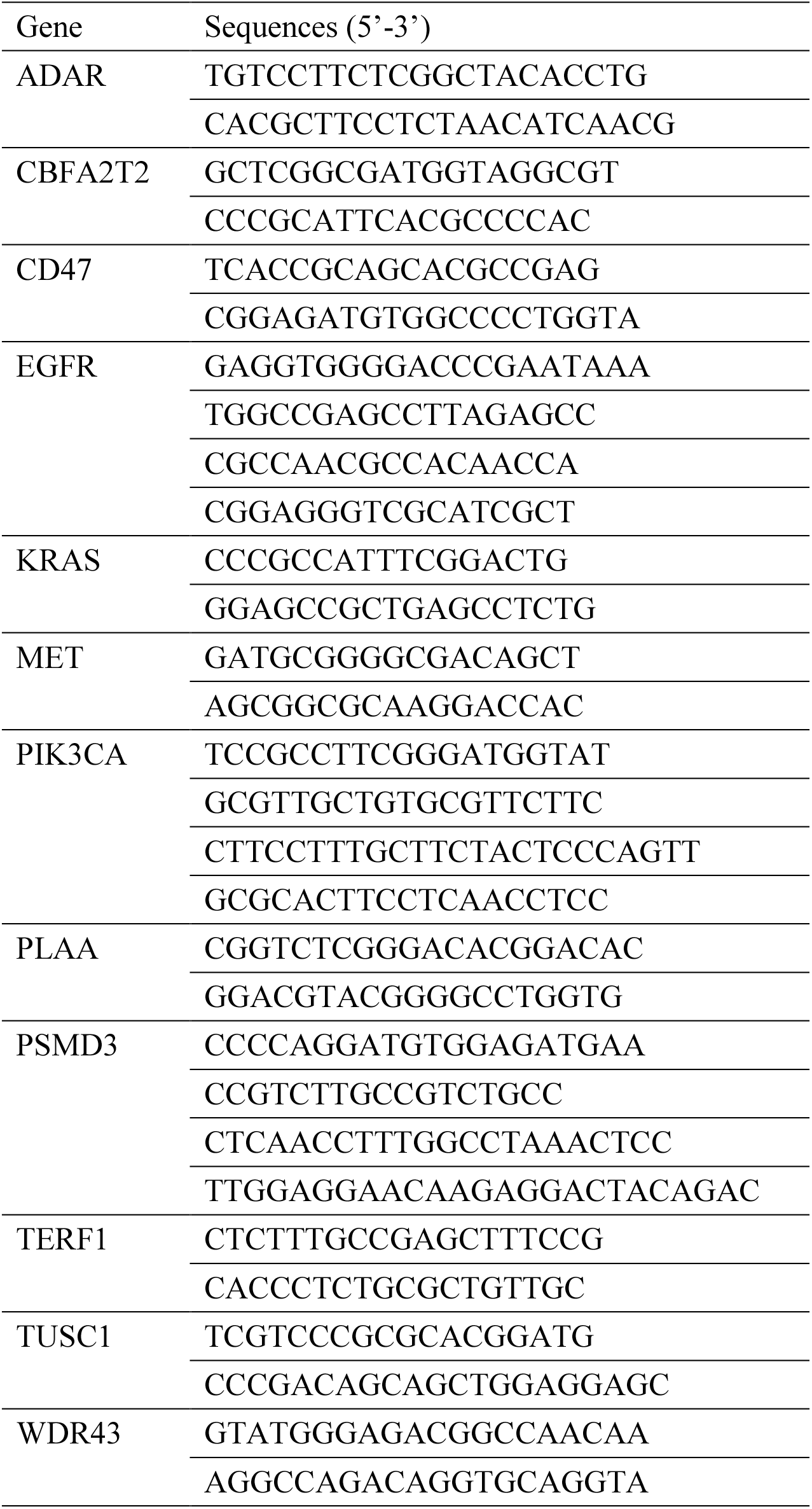
ChIP-qPCR primers.

